# asmbPLS: Adaptive Sparse Multi-block Partial Least Square for Survival Prediction using Multi-Omics Data

**DOI:** 10.1101/2023.04.03.535442

**Authors:** Runzhi Zhang, Susmita Datta

## Abstract

**Background:** As high-throughput studies advance, more and more high-dimensional multi-omics data are available and collected from the same patient cohort. Using multi-omics data as predictors to predict survival outcomes is challenging due to the complex structure of such data.

**Results:** In this article, we introduce an adaptive sparse multi-block partial least square (asmbPLS) regression method by assigning different penalty factors to different blocks in different PLS components for feature selection and prediction. We compared the proposed method with several competitive algorithms in many aspects including prediction performance, feature selection and computation efficiency. The performance and the efficiency of our method were demonstrated using both the simulated and the real data.

**Conclusions:** In summary, asmbPLS achieved a competitive performance in prediction, feature selection, and computation efficiency. We anticipate asmbPLS to be a valuable tool for multi-omics research. An R package called *asmbPLS* implementing this method is made publicly available on GitHub.

## Background

The high-throughput technology has experienced a tremendous improvement, which enables the generation of data on a large scale (omics data) quickly and cost-effectively. Nowadays, omics data from various disciplines such as genomics, transcriptomics, epigenomics, proteomics, microbiomics, and metabolomics [1] have enriched our understanding of the mechanism of different diseases. For instance, molecular phenotyping using genetic and genomic information will allow early and more accurate prediction and diagnosis of disease and of disease progression [2]. In addition, microbiome-derived metabolites have been identified as contributing to a wide range of diseases such as inflammatory bowel disease [3], colorectal cancer [4], type II diabetes [5], asthma [6], as well as obesity [7].

In the past decade, many bioinformatics tools have been developed to enable the analysis of individual omics data [8-10]. With the development of high-throughput studies, there exist more and more patients with different types of omics data available for the same patient. Therefore, the researchers’ interests have gradually moved from single-omics analysis to multi-omics analysis. Since each type of omics data may contribute unique information, the integration of multi-omics data enables us to understand the complex relationship between the omics data and the host systematically and holistically. Furthermore, it helps us build prediction models with good accuracy and then predict outcomes, such as the survival time of patients. For multi-omics analysis, the data set is structured in blocks, where each block represents variables from a specific type of omics data observed on a group of individuals. Usually, multi-omics data has several important characteristics: 1) high dimensionality, the number of features typically vastly outnumbers sample size; 2) sparsity, some features are detectable only in a minority of samples; 3) there exist interactions between features across different blocks. In addition, variables from different blocks can differ in number and nature.

Until now, there has been a vast amount of literature developed for multi-omics data prediction. Sparse Group Lasso (SGL) [11] is a Lasso-based algorithm, which takes group information into consideration. This method incorporates a convex combination of the standard Lasso penalty and the group-Lasso penalty [12]. Integrative Lasso with Penalty Factors (IPF-Lasso) is also an extension of the standard Lasso [13], which considers the group structure by assigning different penalty factors to different blocks for feature selection and prediction. Priority-Lasso [14] is another Lasso-based method designed for the incorporation of different groups of variables. The aim of the priority-Lasso is to define a priority order for different groups of variables. In addition, Random Forest [15] is another powerful prediction algorithm that is known for its ability to capture complex dependency patterns between the predictors and the outcome. One extension of Random Forest called Block Forest [16] has been developed for the consideration of the group structure.

Partial least squares (PLS) regression [17] has also been used for dimension reduction and prediction. Multi-block PLS (mbPLS) [18, 19], although designed for chemical systems at first, can be used for outcome prediction using multi-omics data since the chemical data shares a similar group structure with multi-omics data. Furthermore, sparse mbPLS (smbPLS) [20] has been developed to apply in the bioinformatics field. In addition, sequential mbPLS [21] was proposed to improve the interpretability of multiblock data structure. In smbPLS algorithm, the penalty factor *λ*^*b*^ is fixed for all the PLS components in block *b* (For more details, please refer to “Adaptive Sparse Multi-block PLS Algorithm” subsection in the Method section). However, when all the weights in block b are smaller than *λ*^*b*^ during the convergence iterations given inappropriate *λ*^*b*^, all the weights will shrink to zero, resulting in no information on block *b* included. Therefore, block *b* cannot provide any information for the prediction. In addition, *λ*^*b*^ is not necessarily the best penalty factor for all the PLS components in block *b*, since in different components there might be different proportions of relevant variables (different numbers of non-zero weights). Motivated by these, we proposed a new multi-omics prediction model named adaptive sparse multi-block partial least square (asmbPLS), which allows different penalty factors for different PLS components of block *b* by selecting the specific quantile of feature weights as the penalty factor. By doing this, we can always obtain information on the relative importance of features for each block. The aim of the proposed method is to obtain the subset of features that are most associated with the outcome and then predict the outcome using the selected features. Also, using the quantile of feature weights as the penalty factor will make the interpretation more straightforward.

For predicting the phenotypic features such as the survival outcome of patients or time till a severe disease development, many methods have been developed using the Cox model [22-24], whose efficiency is severely dependent on the proportional hazard assumption. In occasions where the proportional hazard assumption is violated, the accelerated failure time (AFT) model [25] is an appealing alternative to predict patient survival. In the AFT model, suppose T denotes time to a certain event of interest, we model a transformed *Y* = *log*(*T*) on a set of covariates *X*_1_, …, *X*_P_ using linear regression. In this paper, we implemented asmbPLS to predict the log-transformed survival time. To address censoring in the survival outcome, we implemented mean imputation [26] for the right-censored data. The validity of the method is confirmed using simulated data and real data. Note that the method can be easily applied to different types of omics data, preprocessed depending on the type of data. Also, the outcome used here is not limited to the survival time, any continuous outcome can be fitted into this method. An R package called *asmbPLS* implementing this method is made publicly available on GitHub (https://github.com/RunzhiZ/asmbPLS).

## Results

### Simulation Strategies

We simulate *n* samples, *q* bacterial taxa and *p* metabolites to mimic the real microbiome and metabolome data. And censored survival time is then generated based on the simulated microbiome and metabolome data.

#### Microbiome Data

The microbiome data is simulated using the Dirichlet-multinomial (DM) distribution [27] to model the over-dispersed taxon counts. We denote ***Q*** = (*Q*_1_,*Q*_2_,…,*Q*_*q*_) as the observed counts for *q* bacterial taxa. The most common model for count data is the multinomial model, whose probability function is given as:

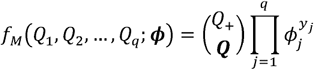

where 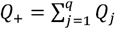 is the total taxon count, which is determined by the sequencing depth, and ***ϕ*** = (*ϕ*_1_, *ϕ*_2_,…, *ϕ*_q_) are underlying taxon proportions with ∑ *ϕ*_j_ = 1. *Q*_+_ can be different for different samples due to the variability of the sequencing depth.

The DM distribution assumes that proportions ***ϕ*** used in the multinomial model come from the Dirichlet distribution [28] and the probability function is given by:

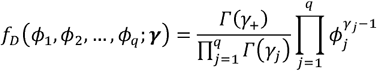

where ***γ*** = (*γ*_1_, *γ*_2_,…, *γ*_*q*_) are positive parameters, which can be generated from the uniform distribution, 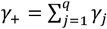 and *Γ*(·) is the Gamma function. The DM distribution then results from a compound multinomial distribution with weights from the Dirichlet distribution:

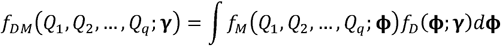

In summary, to generate microbiome data, we first generate ***γ* ∼** *uniform*(0, 1). ***γ*** is then used for generating ***ϕ***, and the generated ***ϕ*** with the *Q*_+∼_*uniform*(m,2m) can be used to generate taxon counts for each sample. Therefore, a *n* × *q* microbiome data matrix ***X***^**micro.*count***^ is generated where rows indicate the samples and columns indicate the microbial taxa. Microbial relative abundance ***X***^**micro.*relative abundance***^ is then calculated from the count data, which will be used in the downstream simulation.

#### Metabolome Data

Once microbiome data are generated, they can be used to generate the metabolome data, since there are associations between microbiome data and metabolome data. In other words, the metabolite levels can depend on the levels of microbial taxa. Notice that ***X*** ^**micro.*relative abundance***^ should be scaled first to control the microbial effect. Let a *n* x *p* matrix ***X***^***meta***^ be the simulated metabolome data, let 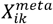 be the intensity of *k*th metabolite in *i*th sample, we assume that the metabolite level of 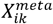 is consisted of three parts:

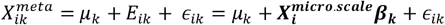

where *μ*_*k*_ denotes the average intensity of metabolite *k*, 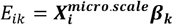 is the microbial effect for metabolite *k* in *i*th sample and *∈*_*ik*_ is the random error term. Here, we have taken *μ*_*k*_ ∼ *uniform*(4, 8) and *∈*_*ik*_ ∼ *N*(0,1). ***β***^(***e***)^ = (***β***_**1**_, …, ***β***_***k***_, …, ***β***_***p***_) is a *q* × *p* matrix to indicate the effect of *q* microbial taxa on *p* metabolites. Among the *q* x *p* pairs in the ***β***^(***e***)^ matrix, *e* pairs will be selected randomly to have non-zero values with all the other elements in ***β***^(***e***)^ to equal to zero. And among these *e* pairs, half of the pairs are randomly selected to have positive values with another half to have negative values. In other words, 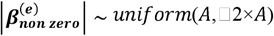, different values of *A* here indicate different scales of association between microbiome and metabolome data.

#### Censored Survival Time

After the generation of the metabolome data, it is scaled also and combined with the scaled microbiome data for survival time generation. Under the AFT setting, we assume that both microbiome data and metabolome data have effects on the true survival time:

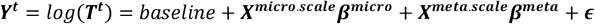

where *baselin* = *log*(500) indicates the logarithm of the baseline survival time for subjects, ***β***^***micro***^ and ***β***^***meta***^ are the coefficients to decide the associations between the features and the survival time, ***∈*** is the random error term. Two types of error distributions are considered here: 1)

Normal distribution, indicating the lognormal distribution for survival time; 2) logarithms of Weibull distribution, resulting in the Weibull distribution for survival time. Specifically, in both cases we take 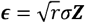, where ***Z*** is either *N*(0, 1) or 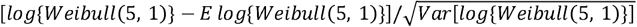, *σ*^2^ = ***β***^***T***^***Σ***_***X***_***β*** with 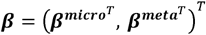 and *r* is the noise to signal ratio. In addition, to simulate the censored time, the censoring variable *c* is taken to be 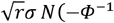, where ***Φ*** is the cumulative distribution function of the standard normal distribution. It will be added to the ***Y*** ^***t***^ to form the ***Y*** ^***c***^ if *c* ≤ 0. For *c* > 0, the true survival time ***Y***^*t*^ will be used.

For ***β*** mentioned above, we considered six settings that we believe cover a broad range of situations. Details are listed in **Table 1**. In setting (1), there are only several relevant features in both blocks. In setting (2), for both blocks, the coefficients are fast decaying with only a small proportion of features contributing to the outcome. Setting (3) is similar to setting (2) but with a slower decay. Setting (4) corresponds to the situation where all the features have equal contributions to the outcome. In setting (5), there are different numbers of relevant features in different blocks (number of relevant taxa > number of relevant metabolites). Setting (6) is similar to setting (5) but with the number of relevant taxa < the number of relevant metabolites. We normalized the vector of ***β*** in each case to control the effect of features on survival and for computational stability.

**Table 1.**
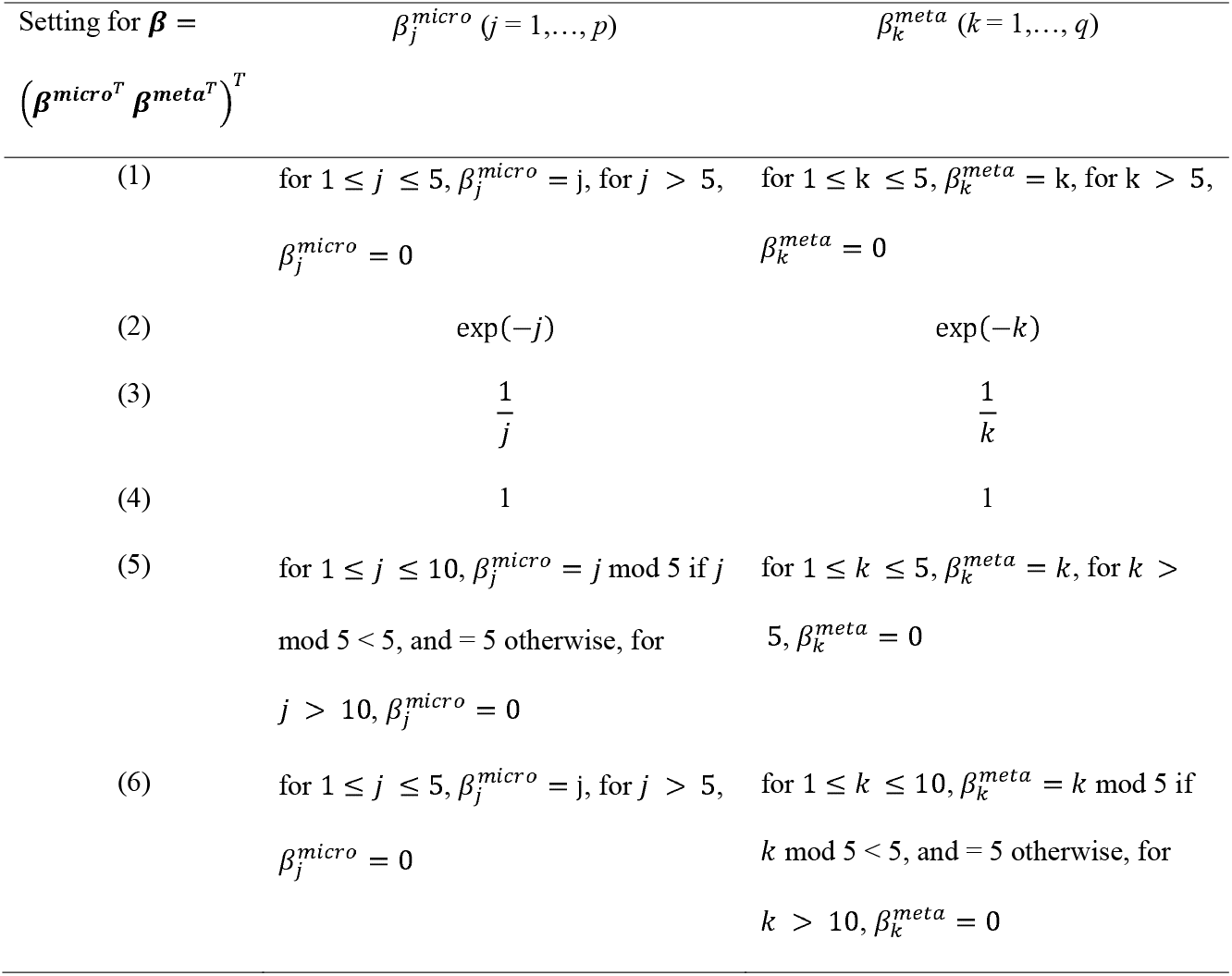
Different settings for **β**. *j* and *k* are the indices for microbial taxa and metabolites, respectively.

### Simulation Results

A variety of simulation settings were considered to account for different scenarios. In low dimension setting, *q* and *p* were taken to be 200, the other parameter values used in this simulation were: *m* = 20,000, *e* = 200, *A* = 0.5 or 2 to denote moderate or high correlation between microbiome and metabolome data, *r* = 0, 0.1, 0.2, 0.5, 1 to indicate different scales of noise and censoring rate (*cr*) = 0.1, 0.3, 0.5, 0.7 to simulate different censoring rates in survival analysis. Sample size *n* was taken to be 100, and an additional *n*_*test*_ = 100 samples were generated using the same design parameters to serve as the test set. For each scenario, we simulated 100 datasets. We also conducted simulations in mixed dimension (*q* = 1,000 and *p* = 200, *e* = 500) and high dimension (*q* = *p* = 1,000, *e* = 1,200) settings, the results are similar to what we present below. In addition to the comparison between different methods, we have also conducted asmbPLS with different pre-defined quantile combinations to see how the performance of asmbPLS is affected by the quantile combinations used. More details can be found in Additional file 1.

Since the microbial data were presented in the form of relative abundance, we imputed the zero value with the small pseudo value 0.5 and then implemented centered log-ratio transformation [29] to the relative data. We conducted the same procedure for the real data. Considering the long computation time of SGL, we removed it in the high dimension setting.

The numbers of PLS components used for asmbPLS and mbPLS were both taken to be 5 in the simulation study. For asmbPLS, the quantile combinations used for cross-validation (CV) in low dimension setting were 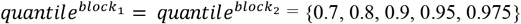 the quantile combinations used for CV in the mixed dimension setting were 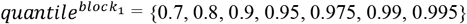 and 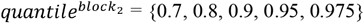 and the quantile combinations used for CV in high dimension setting were 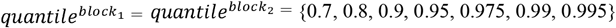. For IPF-Lasso, 9 different penalty ratios were used, i.e. {1:1, 2:1, 1:2, 3:1, 1:3, 4:1, 1:4, 5:1, 1:5}, which indicates different penalty factors used for two predictor blocks.

#### Prediction Performance

The performances of prediction of all the methods were measured in terms of the scaled MSE using the test set:

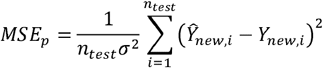

**Figure 1** presents the distribution of the lowest *MSE*_*p*_ using different numbers of PLS components in the low dimension setting. Using the first PLS component or the first two PLS components gives us the lowest *MSE*_*p*_ in scenarios with lower noise (*r* = 0, 0.1, 0.2) most of the time. And with the increase in noise, more than half of the lowest MSEs are obtained from using the first PLS component only. We found a similar trend in other dimension settings also. To comprehensively investigate the performance of asmbPLS, we included asmbPLS with the first PLS component also for comparison.

**Figure 1.**
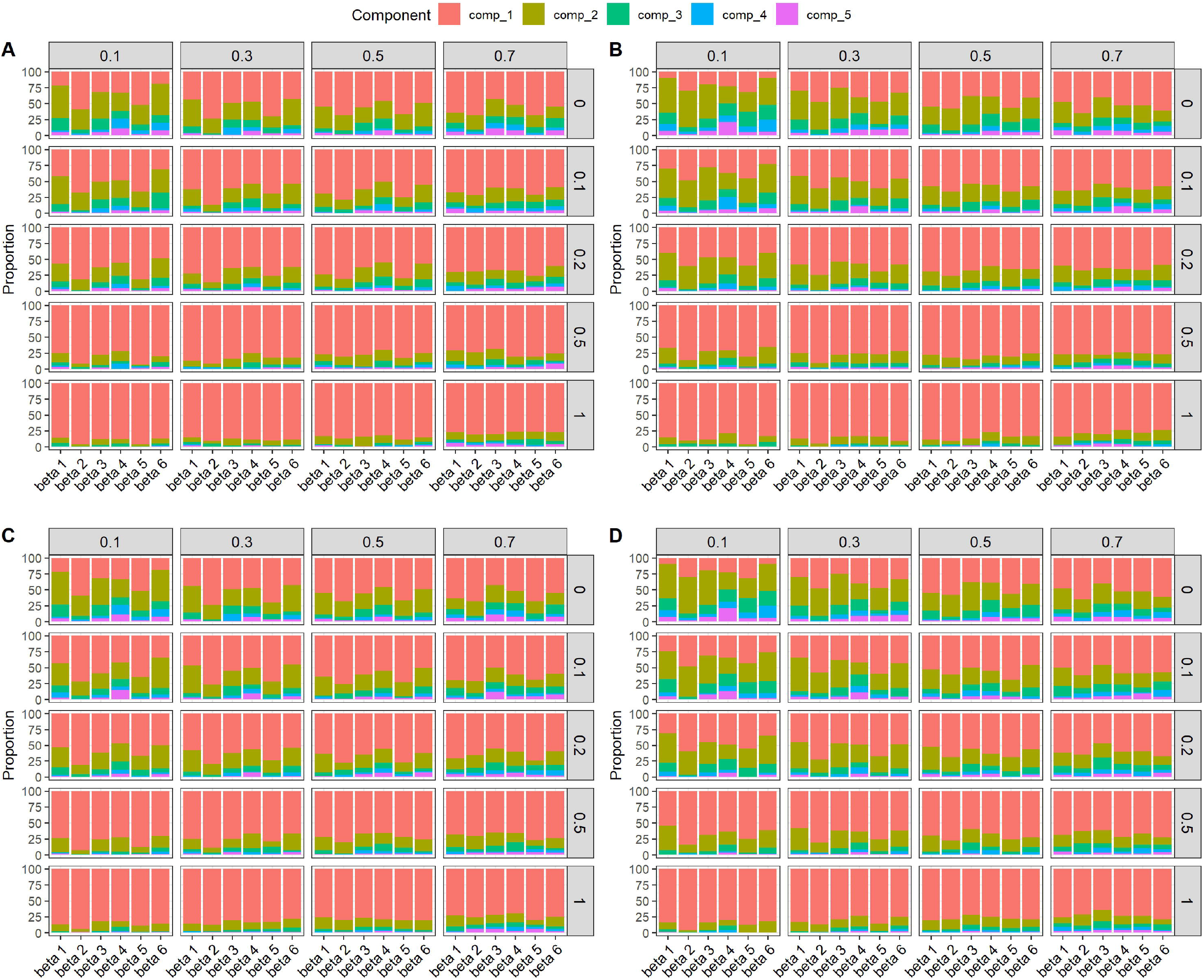
The distribution of the lowest *MSE*_*p*_ using different numbers of PLS components in the low dimension setting. **(A)** Lognormal distributed survival time, ***A*** = 2; **(B)** Lognormal distributed survival time, ***A*** = 0.5; **(C)** Weibull distributed survival time, ***A*** = 2; **(D)** Weibull distributed survival time, ***A*** = 0.5.

**Figure 2** displays the results for low dimension setting with lognormal distributed survival time and *A* = 2. The results of Priority-Lasso were not included here due to its much worse performance. For *cr* = 0.1 (**Figure 2.A**), IPF-Lasso performs the best in most of the scenarios except β setting (4), where all the features have equal contributions to the outcome. In β setting (4), mbPLS, which can be considered as a special case of asmbPLS, performs the best, it makes sense since mbPLS makes use of the information from all the features. For *cr* = 0.3 (**Figure 2.B**), the scaled MSEs of all the methods increase due to the increase in *cr*. IPF-Lasso still performs the best in most of the scenarios except βsetting (4), however, the performance of asmbPLS is closer to IPF-Lasso in all the scenarios. For *cr* = 0.5 (**Figure 2.C**), in scenarios with less relevant features such as βsetting (2), asmbPLS performs the best and slightly better than IPF-Lasso and SGL. In scenarios with more relevant features such as β settings (1)(3)(5)(6), the overall performance of asmbPLS is close to the best. For *cr* = 0.7 (**Figure 2.D**), asmbPLS exhibited better performance than other methods in most of the scenarios except the settings with high noise (*r* = 1), where Block Forest performs the best. In summary, although asmbPLS is not the best in low censoring rate scenarios, the performance of asmbPLS improves with the increase of the censoring rate, and asmbPLS achieves the best performance in high censoring rate scenarios. In addition, asmbPLS performs better in scenarios with less significant relevant features, i.e. *β* setting (2), than the other scenarios. Furthermore, asmbPLS cannot always keep the best performance with the increase in noise.

**Figure 2.**
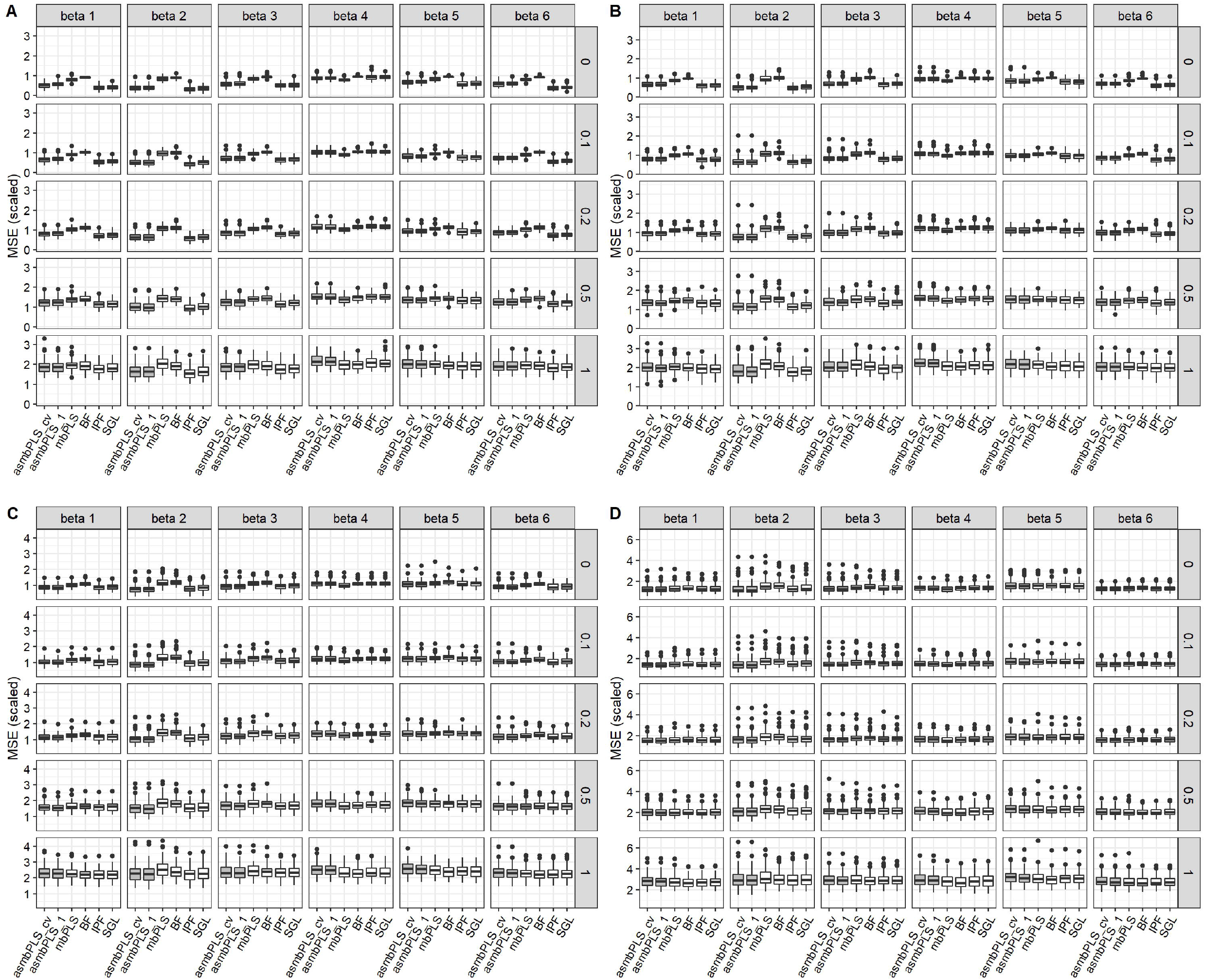
Prediction results for low dimension setting with lognormal distributed survival time and ***A*** = 2. **(A) *cr*** = 0.1; **(B) *cr*** = 0.3; **(C) *cr*** = 0.5; **(D) *cr*** = 0.7.

And by comparing two asmbPLS algorithms with different strategies for selecting the number of components, i.e. asmbPLS_cv (selection of the number of components based on the results of cross validation (CV)) and asmbPLS_1 (only the first component selected), asmbPLS_cv performs better in scenarios with lower censoring rates and lower noise but slightly worse with the increase of the censoring rate and noise.

Similar trends can be found in low dimension setting with lognormal survival time and *A* = 0.5 (Additional file 1: **Figure S1**). In addition, we found similar results also when the distribution of survival time was changed from lognormal to Weibull (Additional file 1: **Figure S2** and **S3**).

#### Feature Selection

In this section, only results for *β* settings (1)(2)(3)(5)(6) are presented since the feature selection for *β* setting (4) is not necessary. In addition, among all the methods, the feature selection nprocedure is not included in mbPLS and Block Forest. Therefore, we compared asmbPLS with IPF-Lasso, Priority-Lasso, and SGL only. The performances of feature selection were measured in terms of sensitivity and specificity.

For different *β* settings, we defined the true relevant features differently. For *β* settings (1)(5)(6), we defined the true relevant features as the features with positive *β* For *β* settings (2)(3), since all *β*s are positive with different values, we defined the true relevant features as the features with false discovery rate (FDR) adjusted p-value < 0.05. After we obtained all the feature selection results, we calculated the true positive count, true negative count, false positive count, false negative count, and further sensitivity, and specificity based on 100 simulated datasets for each scenario.

**Figure 3** presents the sensitivity and specificity of the feature selection for low dimension setting with lognormal distributed survival time. According to **Figure 3**, the performances of different methods on feature selection vary in different types of omics data and all methods show a similar trend in different *A* settings. Regarding the sensitivity (**Figure 3.A** and **3.C**), the performance of asmbPLS might not be the best at lower noise with *β* settings (1)(5)(6) in scenarios with *cr* = 0.1, 0.3 or 0.5. However, with the increase of the noise, asmbPLS gradually performs better than the other methods whose sensitivity is largely affected by the noise. For *β* settings (2)(3), asmbPLS performs the best with sensitivity close to 1 in all the scenarios regardless of the censoring rate and the noise. This can be due to the definition of the true relevant features in these two *β* settings because the nature of asmbPLS makes it always find the most significant features (with the lowest p-value). For *β* settings (1)(5)(6), the true relevant features are defined as the features with positive *β*. However, sometimes the features with positive β are not the significant features in the simulations, which causes lower sensitivity in these three settings for all the methods. In scenarios with high censoring rates (0.7), the sensitivity of asmbPLS is the highest among all the methods, which is corresponding to the lower MSE of asmbPLS. Correspondingly, the better sensitivity of asmbPLS is along with the lower specificity (**Figure 3.B** and **3.D**). It is worth noting that the specificity of other methods increases with the increase of noise. On the contrary, there has been a gradual decline in the specificity of asmbPLS. In other words, asmbPLS tends to retain more features than other methods, especially in scenarios with higher noise, which corresponds to the reduced prediction performance of asmbPLS with the increase in noise since including more irrelevant features makes the prediction more difficult. For asmbPLS, although with more features included and more false positive counts, the weights are assigned differently for the different blocks and features, which can still help us to find the most significant features. We found similar results with Weibull distributed survival time (Additional file 1: **Figure S4**).

**Figure 3.**
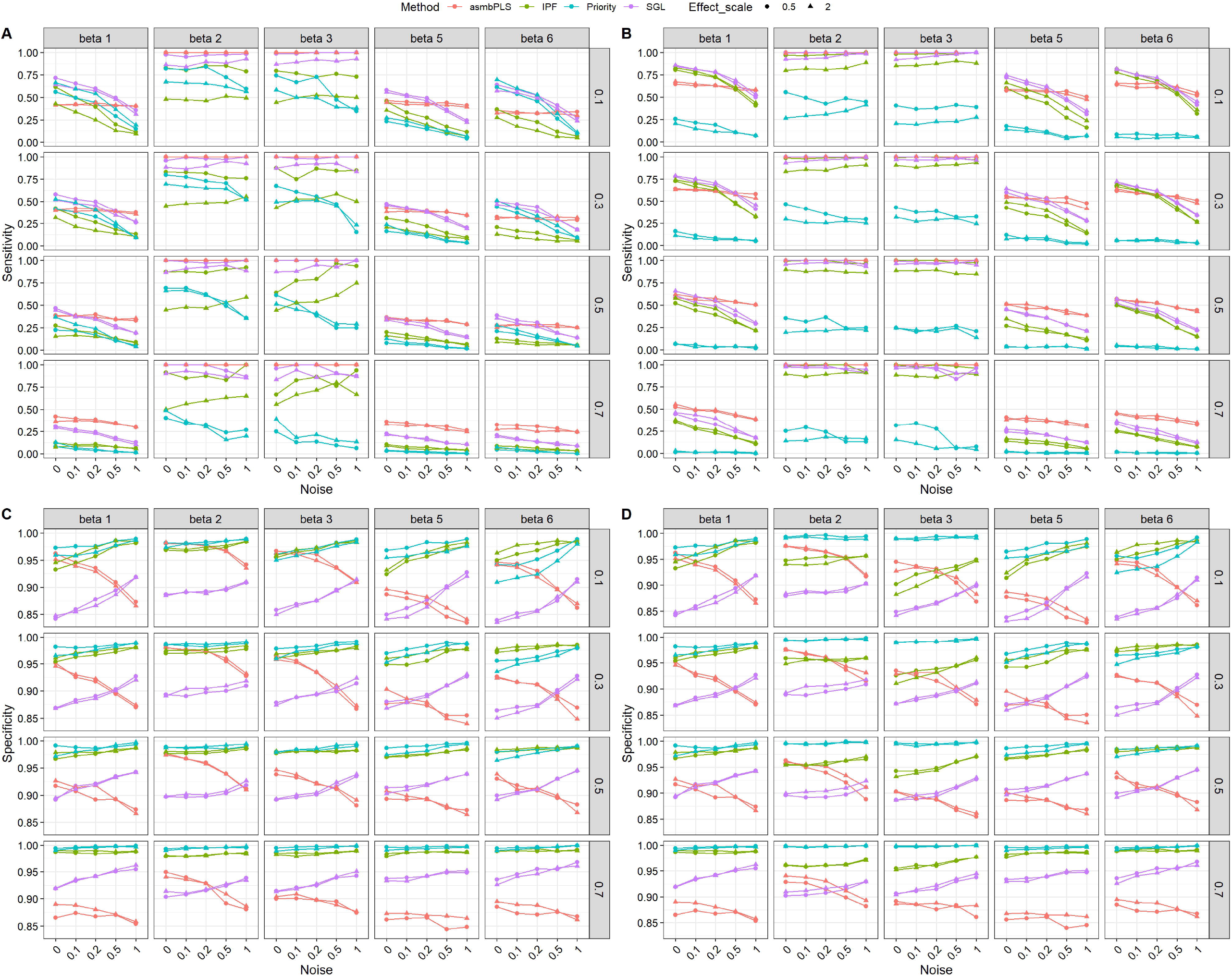
Sensitivity and specificity of the feature selection for low dimension setting with lognormal distributed survival time. **(A)** Sensitivity for microbiome block; **(B)** Sensitivity for metabolome block; **(C)** Specificity for microbiome block; **(D)** Specificity for metabolome block.

#### Computation Efficiency

**Table 2** displays the mean computation time for each method in different dimension settings. The computation time is measured as the time needed for model fitting including the CV. Among all the methods, mbPLS, which does not require the CV step, is the fastest procedure, followed by Priority-Lasso and asmbPLS. As the special case of asmbPLS, the computation time of mbPLS is equal to the computation time of asmbPLS given selected quantile combinations for all the PLS components. For asmbPLS, the CV step takes the longest time, which largely depends on the number of quantile combinations and the number of folds for the CV. Overall, asmbPLS algorithm is faster than IPF-Lasso, Block Forest and SGL, especially when compared to Block Forest, and SGL. One thing to notice is that IPF-Lasso shows a decreased computation time in the mixed dimension and high dimension settings compared to low dimension setting. The computation time of all the other methods increase in different degrees with the increases of the dimension. SGL is the slowest among all the methods, which is not implemented in the high dimension setting for simplicity.

**Table 2.** Average computation time (in seconds) for different methods in different dimension settings.

| Dimension | asmbPLS_0.5 | asmbPLS_0.7 | asmbPLS_0.9 | mbPLS | BF | IPF | SGL | Priority |
| --- | --- | --- | --- | --- | --- | --- | --- | --- |
| Low | 3.17 | 1.63 | 1.03 | 0.0018 | 50.04 | 8.11 | 83.51 | 0.57 |
| Mixed | 6.18 | 3.38 | 2.32 | 0.0047 | 73.32 | 6.12 | 226.78 | 0.53 |
| High | 11.25 | 6.79 | 4.98 | 0.0072 | 101.54 | 7.81 | - | 0.51 |

### Application to Melanoma Patients Data

We used data from [30], which includes progression-free survival (PFS) interval/status, clinical variables, and various types of omics data collected for 167 melanoma patients. Among those collected in this study, we consider three blocks, one low-dimensional block (clinical covariates) and two high-dimensional blocks, i.e. microbiome data and proteomics data. The outcome is progression-free survival interval, the patients with the unobserved event (patients with no detection of disease progression or death) are considered right-censored.

Among the available clinical variables, we selected 17 clinical variables, age and BMI are continuous with all the other variables are binary: sex, tumor response status, types of treatment, substage of disease in patients with late-stage disease, dietary fiber intake level, lactate dehydrogenase (LDH) level, probiotics use at baseline, antibiotic use at baseline, metformin use at baseline, steroid use at baseline, statin use at baseline, proton pump inhibitor (PPI) use at baseline, beta-blocker use at baseline, other (non beta-blocker) hypertensive medication use at baseline, whether or not the patient received system therapy prior to baseline. The microbiome block and the proteomics block contain 225 microbial features and 6773 proteins, respectively. After filtering the subjects with missing values and without survival data, we obtained 89 samples (55 events) for the downstream implementation. The mean imputation was conducted first for the censored survival time, we then applied the AFT model to determine whether there is an association between PFS and any of the features in the three blocks. Since we have no prior information about the association, we implemented a univariate analysis for each feature to determine whether any of the features are predictive of PFS. To this end, we fitted the simple linear regression model with imputed log-survival PFS as our outcome and each feature as our predictor one at a time. The obtained p-values in each block were then adjusted using the false discovery rate (FDR). Specifically, at a significance level of 5%, no microbial features and proteins are significant. And for clinical variables, only tumor response status and substage of disease are significant. With this information, we applied asmbPLS and the other methods in the real data and compared them in three aspects: 1) The fit of the model to data; 2) Prediction error of the model; 3) Feature selection by the model.

To measure the fit of the model to data, we used the MSE of fit to compare different methods:

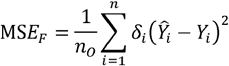

where *n*_*o*_ is the number of observed samples. We included the first three PLS components for comparison for PLS-based methods. In addition, for asmbPLS, the 5-fold CV was implemented to obtain the best quantile combination used for model fitting. The pre-determined quantile combinations set were *quantile*^*microbiome*^ = {0.9, 0.925, 0.95, 0.975, 0.99, 0.999}, *quantile* ^*proteomics*^ = {0.997, 0.9985, 0.9993, 0.9999}, *quantile* ^*clinical*^ = {0, 0.3, 0.5, 0.7, 0.8, 0.9, 0.99}, resulting in 168 combinations considered. Based on the results of CV, the optimal number of PLS components is 2, combination (0.975, 0.997, 0.9) was selected for the first PLS component, and combination (0.999, 0.9999, 0.9) was selected for the second PLS component. **Table 3** lists the values of *MSE*_*F*_. We can see that for both PLS-based methods, including one more PLS component decreases the *MSE*_*F*_, which is always the case for PLS model fitting. Among all the methods, mbPLS with 2 PLS components shows the lowest MSE since this non-sparse method uses all the features for fitting, which could result in over-fitting. For other methods, asmbPLS with 2 PLS components performs a little bit better than SGL and IPF-Lasso and much better than Block Forest. The performance of Priority-Lasso is much worse than all the other methods, which is corresponding to the results from the simulation study.

**Table 3.** Comparison of the model fitting and prediction performance for different methods using the real data.

| Methods | $MSE_F$ | $MSE_{P,LOO}$ |
| --- | --- | --- |
| asmbPLS with 1 PLS component | 0.776 | 0.820 |
| asmbPLS with 2 PLS components | 0.674 | 0.764 |
| mbPLS with 1 PLS component | 0.834 | 1.426 |
| mbPLS with 2 PLS components | 0.424 | 1.149 |
| Block Forest | 1.006 | - |
| IPF-Lasso | 0.740 | 0.804 |
| Sparse Group Lasso | 0.698 | 0.778 |
| Priority-Lasso | 12.303 | 24.468 |

On the other hand, to evaluate the prediction performance of the model, we computed the MSE of prediction via leave-one-out (LOO) CV with each observed sample serving as validation data for once:

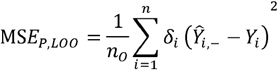

where *Ŷ*_*i*,-_ is computed by the model using the dataset with *i*th sample excluded. The selected quantile combination from the previous CV was used here for the validation set. The results are listed in **Table 3**. Block Forest was able to fit the model, but cannot be used to do the prediction due to the error message in R. As seen in the table, among all the other methods, asmbPLS with 2 PLS components performed the best, followed by SGL, IPF-Lasso, mbPLS, and Priority-Lasso. Even though the performance of asmbPLS is only slightly better than SGL, asmbPLS can still be a better choice considering the relatively much less computation time.

For asmbPLS, 6 microbial taxa, 21 proteins, and 2 clinical variables were selected. And block weights for microbiome block, proteomics block, and clinical blocks were 0.117, 0.065, and 0.991, respectively. We found that although some of the features from the microbiome and proteomics blocks were selected, the weights for these two blocks were much smaller compared to the weight of the clinical block, indicating that the variables from the clinical features might be more relevant to the outcome, which was also validated by the univariate analysis. The two clinical variables selected by asmbPLS were tumor response status and substage of disease, which were the only two significant clinical variables. In addition, for the selected features which were assigned with high weight, all of them were significant before FDR adjustment. For IPF-Lasso, 0 microbial feature, 1 protein, and 1 clinical variable were selected. The variable substage of disease, which was highly significant before and after FDR adjustment, was not selected by IPF-Lasso. For SGL, 0 microbial feature, 0 protein, and 3 clinical variables were selected. In addition to the two significant variables, lactate dehydrogenase (LDH) level, which was not significant before FDR adjustment, was selected and assigned with a much higher weight than substage of disease. For priority-Lasso, no feature was selected.

In addition to the three aspects discussed above. We explored the utility of the super score, which contained the information from all the blocks and can only be obtained from asmbPLS and mbPLS, as the predictor for the outcome. Specifically, an optimal cut-point on super score was determined to define the two groups using the maximally selected rank statistics [31] as implemented in the R package *survminer* [32], and the p-value was calculated based on a log-rank test between the resulting groups. As seen in **Figure 4**, super score is significantly associated with progression-free survival time. Patients with higher super scores (blue group) seem to have much higher survival probability. In other words, once we have a new sample with its corresponding microbiome, proteomics, and clinical data, asmbPLS is able to calculate the super score and then assign the sample to the high or low survival group. This information may help the decision of future treatment.

**Figure 4.**
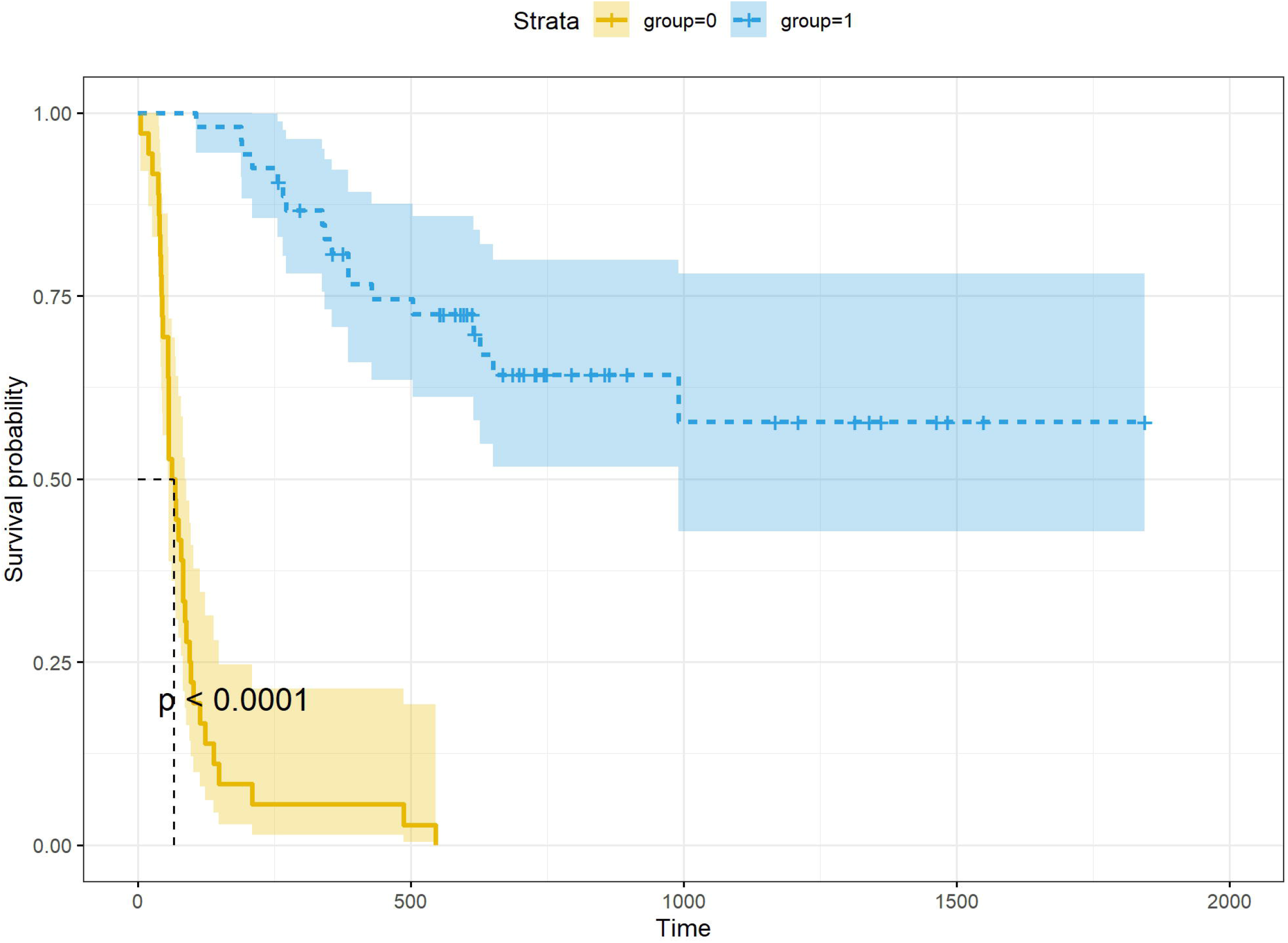
Prediction of PFS using super score from asmbPLS.

Furthermore, we have done the additional analysis without using the most significant clinical block. In this test, only microbiome block and proteomics are included as predictors. The same pre-determined quantile combinations set are used for microbiome and proteomics blocks. Based on the results of CV, the optimal number of PLS components is 1, combination (0.999, 0.9999) was selected for the first PLS component, indicating that 1 microbial taxon and 1 protein were selected. These two features are still the most significant feature in each block even though they are not significant after p-value adjustment. For the other three Lasso-based methods, 0 microbial taxon and 0 protein was selected. **Table 4** lists the results for comparison of model fit and CV. With the fact that no predictor is relevant, all the methods show relatively higher *MSE*_*P,LOO*_ than in **Table 3**. Especially, asmbPLS shows higher *MSE*_*P,LOO*_ than the other Lasso-based methods, which is due to the nature of asmbPLS, which always keeps at least one most relevant predictor for each block. This nature will help the prediction when there are significant predictors but can make the prediction worse if there is no real relevant predictor.

**Table 4.** Comparison of the model fitting and prediction performance for different methods using the real data (without clinical block).

| Methods | $MSE_F$ | $MSE_{P,LOO}$ |
| --- | --- | --- |
| asmbPLS with 1 PLS component | 2.14 | 2.92 |
| mbPLS with 1 PLS component | 1.37 | 3.45 |
| mbPLS with 2 PLS components | 0.87 | 3.05 |
| Block Forest | 2.32 | - |
| IPF-Lasso | 2.21 | 2.28 |
| Sparse Group Lasso | 2.21 | 2.35 |
| Priority-Lasso | 2.21 | 2.26 |

## Discussion and Conclusion

In this paper, we developed asmbPLS algorithm to identify the most significant features of the multi-omics data on the same set of samples and then use the selected features to predict the outcome. Different from smbPLS, asmbPLS is flexible in determining the penalty factor for different omics data in different PLS components. With some prior knowledge of the omics data, the pre-decided quantile set can be provided to each block, and then the best quantile combination can be chosen in a completely data-driven manner. In addition, using the quantile makes the interpretation more straightforward, block with selected quantile = 0.95 indicates that only the top 5% features are relevant to the outcome. asmbPLS works with continuous predictor variables and continuous outcomes, binary variables can be transformed to 0/1 to meet the requirement. And for categorical variables with more than 2 levels, the one-hot encoding can be one strategy, where the categorical variable with G levels can be transformed into G-1 dummy variables. asmbPLS is implemented in the R package *asmbPLS* available on our GitHub and will soon be uploaded to CRAN.

Simulation studies have demonstrated that asmbPLS exhibits better prediction performance than the other methods in scenarios with higher censoring rates, especially where we have less relevant features. According to the results of the simulation study, we found that 5 PLS components may not be necessary considering both the prediction performance and the efficiency since including more PLS components did not stably improve the prediction and requested more computation time in the CV procedure. However, we suggest users always include several more PLS components and then select the optimal number of components based on the results of the CV. In addition, in scenarios with higher censoring rate and noise, asmbPLS with the first component can be a safe choice. Regarding the results of feature selection, asmbPLS is more likely to select the true relevant features than the other methods with the increase in noise, especially in scenarios with higher censoring rates. Due to the nature of asmbPLS, the significant features are always selected, which is corresponding to the results of β settings (2)(3), where the true relevant features are defined as adjusted p-value < 0.05. Although asmbPLS tends to select more features than other methods, the weights assigned by asmbPLS for the features and for the blocks are still informative enough to find the most significant features. The performance of asmbPLS is further validated in the application to real data. In addition, the exploration of the super score in real data has proved the super score of asmbPLS as the predictor for classifying the different survival groups.

One limitation of asmbPLS is that when there is no real relevant feature in the predictor blocks, including at least some features in each block will sacrifice the prediction performance of asmbPLS, even though this will provide more information about the relative importance of the features.

In summary, asmbPLS achieved a competitive performance in prediction, feature selection, and computation efficiency. We anticipate asmbPLS to be a valuable tool for multi-omics research.

## Methods

### Right Censored Data Imputation

In survival analysis, a subject is considered to be censored if time-to-event information is incomplete. Among the different types of censoring, right-censoring is the most common one. Let 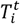 and 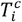 denote the true survival time and censoring time for *i*th subject (*i* = 1, …, *n*), respectively. Then the observed survival time and the event indicator are defined as 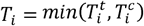 and 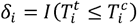, respectively. And observed data for *i*th subject can be presented by (*T*_*i*_, δ_*i*_) where subjects with *δ*_*i*_ = 1 and with *δ*_*i*_ = 0 are called observed and unobserved subjects, respectively. In the AFT model, we consider fitting asmbPLS regression model of *Y*_*i*_ = *log*(*T*_*i*_) on 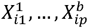, where 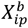 is the *p*th feature of a specific omics data (block *b*) for *i*th sample. Since the 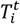 are not available for the individuals with δ_*i*_ = 0, the general prediction method will not apply. In addition, ignoring censored subjects would cause a prediction bias, and reducing sample size also results in loss of power. To handle this, we propose to replace *Y*_*i*_ by 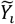, such that (i) it has approximately the same mean function as *Y*_*i*_ and (ii) it is computable from the observed data. In this study, mean imputation [26] is used to this end. Under this scheme, we keep observed *Y*_*i*_ intact but replace unobserved *Y*_*i*_ by its expected value 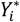 given that the true survival time 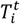 was larger than the censoring time 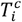. Let *S*(*t*) denotes the survival function, the 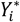 can be estimated from the Kaplan-Meier curve of the survival function: 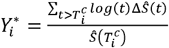, where *Ŝ* is the Kaplan-Meier estimator of the survival time, and Δ*Ŝ*(*t*) is the jump size of *Ŝ* at time t. Note that we need to treat the largest observation as a true failure for this calculation since it is necessary to make a tail correction if the largest observation *t*_*max*_ corresponds to a censored event. In summary, we let 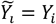 for observed survival time, and 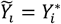 for unobserved survival time for the *i*th sample instead.

### Adaptive Sparse Multi-block PLS Algorithm

In mbPLS, let the input matrix including *B* blocks (different omics data) ***X*** = [***X*** ^**1**^,…, ***X*** ^***b***^,…, ***X*** ^***B***^] and block *Y* be the predictor matrix and outcome vector on the same *n* samples. For *j*th PLS (*j* = 1, 2, …) component, the dimension reduction is implemented by taking a linear combination of the variables to obtain the block score 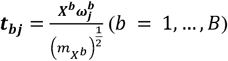 in each block, where 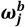 is ***X***^***b***^ block variable weights that express the relevance of variables, which is obtained via the algorithm, and 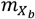 is the number of variables in ***X***^***b***^ block that is used for block scaling. After calculating 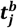 for each block, we combine all block scores in a new matrix 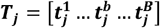, which includes information from different blocks. Then the dimension reduction is conducted again by taking a linear combination of the different block scores to obtain the super score 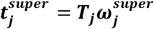, which summarize variable information from different blocks. Similarly, ***u***_***j***_ is a summary vector of ***Y***, i.e. ***u***_***j***_ = ***Yq***_***j***_ (***q***_***j***_ being the ***Y*** weight for *j*th PLS component). The goal nof the mbPLS algorithm is to find the parameters that maximize the covariance between two summary vectors, i.e. 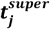 and ***u***_***j***_. Therefore, the problem is formally expressed as follows: 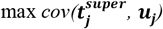 with, 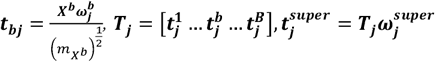 and ***u***_***j***_ =***Yq***_***j***_, subject to 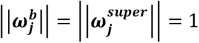.

Once all the parameters are calculated for the first PLS component, ***X*** and ***Y*** are deflated and then the deflated ***X*** and ***Y*** are used for the calculation of the second PLS component and so on. Furthermore, smbPLS [20], a sparse version of the mbPLS, can be achieved by adding an *L*_1_ penalty on each weight vector 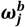. In other words, the variable selection procedure is implemented on each 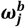, and the weights of unimportant variables will shrink to zero. In smbPLS algorithm, penalty factor *λ*^*b*^ is fixed for all of the PLS components in block *b*. Unlike smbPLS, asmbPLS allows different penalty factors 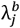 for *j*th PLS component of block *b* by selecting the specific quantile of block weights as the penalty factor. Specifically, each step of the asmbPLS algorithm is listed in **Table 5**. In this algorithm, *quantile* is the function to obtain the corresponding quantile of the absolute weight, and *sparse* is the soft thresholding function *sparsec*(***x***,λ) = *sign*(***x***) (|***x***| − λ)_*+*_ that is used to optimize the objective function with lasso penalties. Using the *quantile* function, we can always find the reasonable 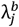 that helps us to retain the most relevant variables. Combining these two functions, we aim to find the best quantile for each PLS component in each block that maximizes the prediction performance of the proposed method. Furthermore, by doing this, the explanation for the retained variables can be more straightforward.

**Table 5.**
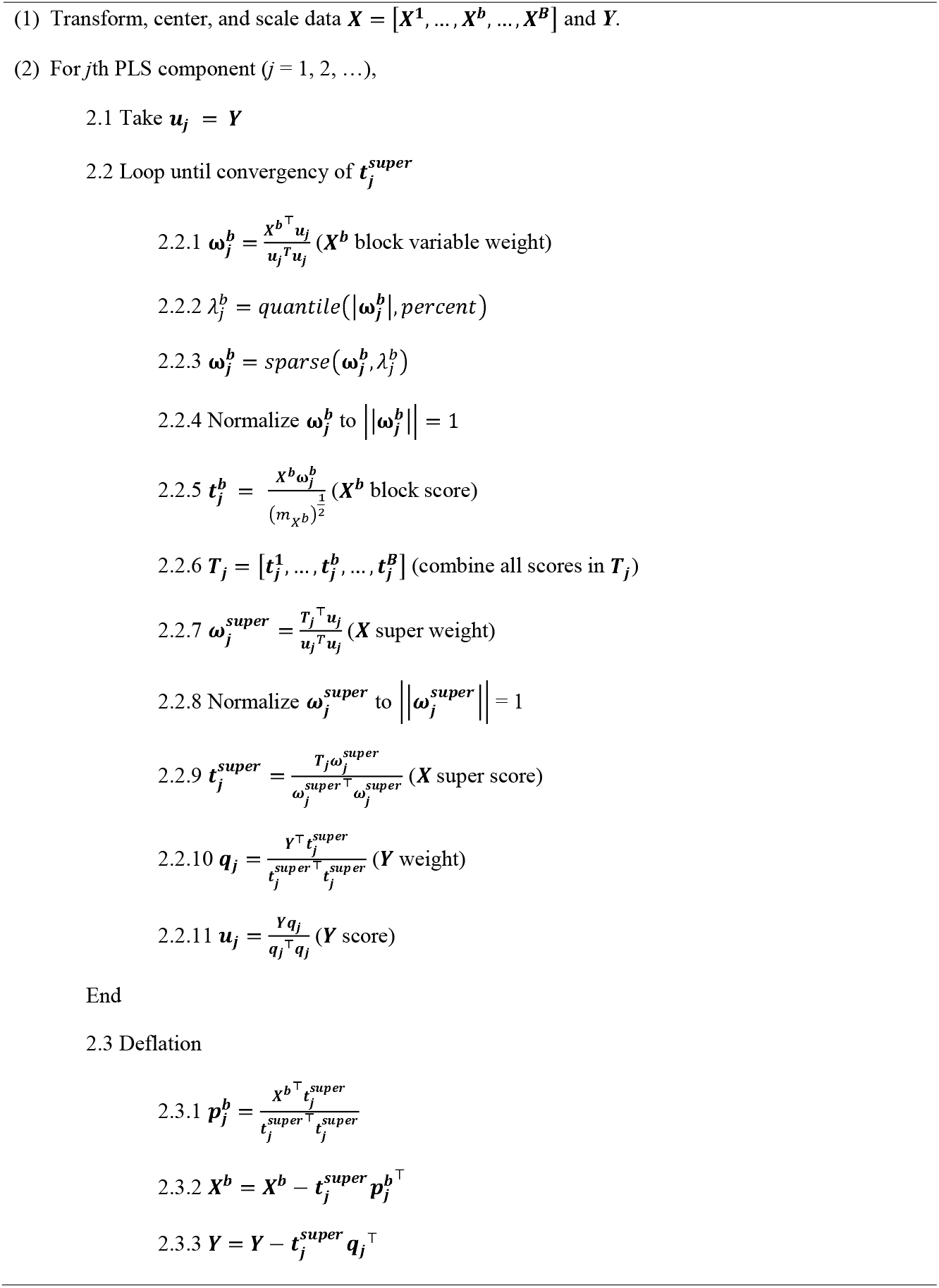
Pseudocode for the asmbPLS algorithm.

After the model fitting, the parameters 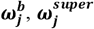, and ***q***_***j***_ could be saved for the prediction when we have the new data ***x***^***new***^. Specifically, 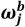 is used to calculate the corresponding 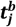 for each block in the new data. And then 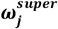 is used for calculating the 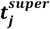 with these Calculated 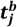. After that, 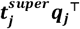 is calculated, which could be used as our prediction based on the first PLS component. If we want to use more PLS components, 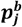 is calculated for obtaining the deflated ***X***^***new***^, which could be further used for calculating 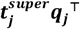 for the second PLS component, and so on. After the 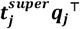 for all the PLS com+ponents are calculated, they can be combined for obtaining the predicted ***Y***. Notice that the mean and standard deviation of each feature of ***X*** and the outcome ***Y*** in the training set will also be saved, which can be used for the scaling of the new ***X*** and obtaining the original scale of the predicted ***Y***.

### Parameter Tuning and Model Selection

We let *B* = 2 and the number of PLS components = 5 here for instance. The CV is used to tune the quantile combination for different PLS components in different blocks. Tuning these parameters is equivalent to choosing the “degree of sparsity”, i.e. the number of non-zero weights for each PLS component in each block. And the chosen quantile combination is the one giving the best prediction. The CV procedure is presented in **Table 6**.

**Table 6.**
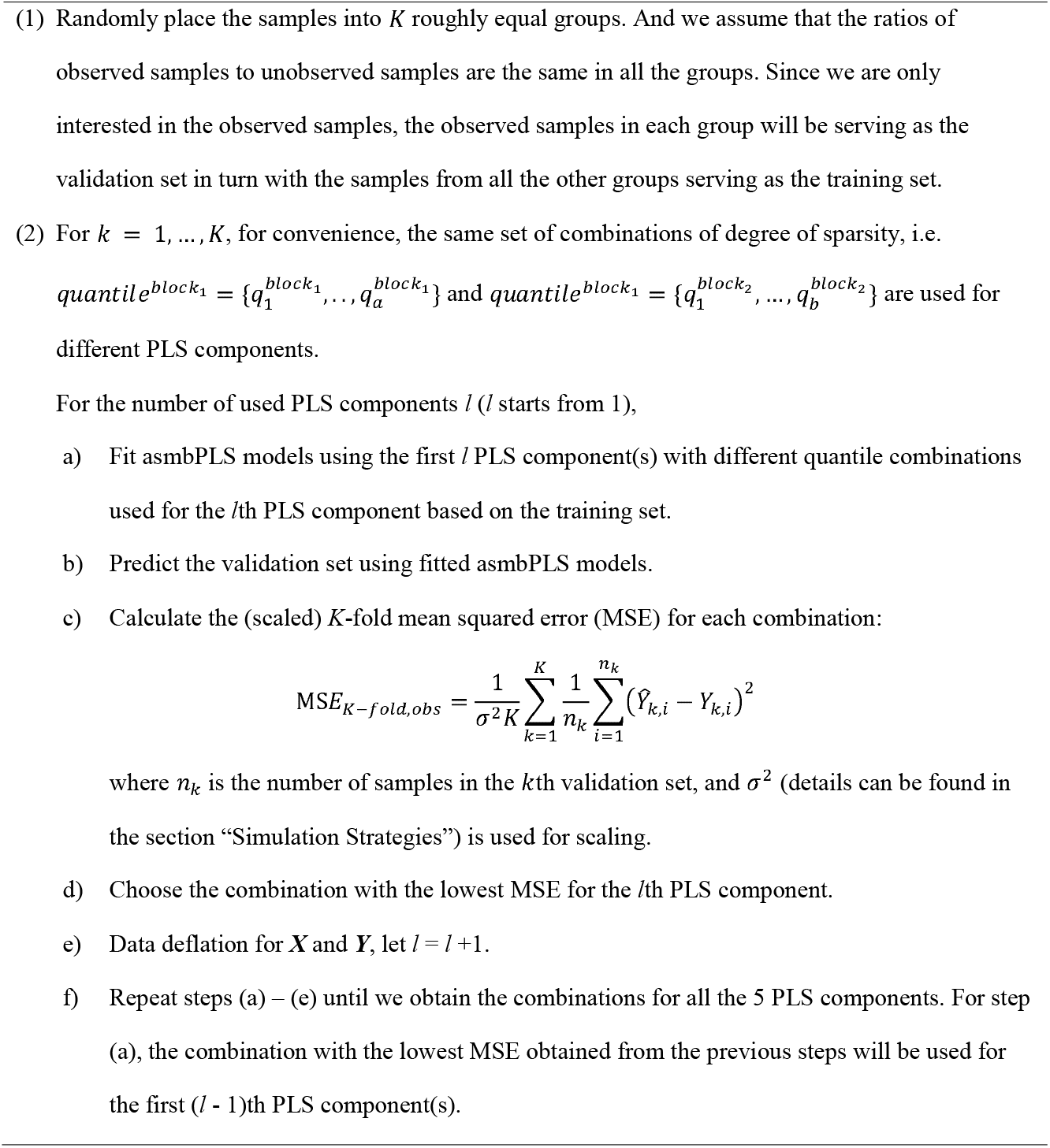
CV procedure for the asmbPLS algorithm.

The selection of *K* and the number of quantile combinations will largely impact the computational efficiency. We choose *K* to be 5 in our study, which is large enough for parameter tuning. The selection of the quantiles for each block should be based on prior knowledge of the corresponding omics data. For example, assuming only a small proportion of features are relevant, then a higher quantile should be considered to retain fewer features.

With the information from the CV, we can determine the number of PLS components used for prediction also. Usually, the optimal number of PLS components is the one that corresponds to the lowest MSE in CV. However, due to the over-fitting issue, we allow the selection of fewer components if the decrease of MSE is slight when one more component is included. The strategy for selecting the number of PLS components is summarized: 1) Let the initial number of components be *comp* = 1; 2) Check whether including one more component decreases the MSE by 5%, i.e. *MSE*_*comp*+1_ ≤ *MSE*_*comp*_ × 0.95; 3) If so, let *comp* = *comp* + 1 and back to step 2), otherwise, let *comp* be the selected number of components.

### Technicalities and Implementation

All the implementations were conducted using R 4.1.0 [33] in the “HiperGator 3.0” high-performance computing cluster, which includes 70,320 cores with 8 GB of RAM on average for each core, at the University of Florida. We compared the proposed method, i.e. asmbPLS, with mbPLS, Block Forest, IPF-Lasso, Sparse Group Lasso, and Priority-Lasso using both the simulated and the real data. The choice of parameters for each method followed the suggestion from the corresponding package tutorial. All these methods make use of group structure information and enable us to do the prediction.

### Data Source

The real data were obtained from [30], which include progression-free survival interval/status, clinical covariates, and various types of omics data such as microbiome data and proteomics data for melanoma patients. The omics data were pre-processed and are presented in the form of relative abundance and parts per million.

## Supporting information

Supplemental materials including the additional data analysis, supplemental tables, and supplemental figures.

## List of abbreviations

*SGL*: *Sparse Group Lasso*
*IPF-Lasso*: Integrative Lasso with Penalty Factors
*PLS*: Partial Lease Square
*mbPLS*: Multi-block Partial Lease Square
*smbPLS*: Sparse Multi-block Partial Lease Square
*asmbPLS*: Adaptive Sparse Multi-block Partial Lease Square
*AFT*: Accelerated Failure Time
*DM*: Dirichlet-multinomial
*CV*: Cross Validation
*MSE*: Mean Squared Error
*FDR*: False Discovery Rate
*PFS*: Progression Free Survival

## Declarations

## Acknowledgments

The authors thank the editor, and the reviewer for their helpful comments and suggestions.

## Author contributions

SD and RZ designed the study, SD provided theoretical support when required, RZ implemented the simulation and the analyses, RZ wrote the manuscript. All the authors have read and approved the final manuscript.

## Availability of data and materials

The data that support the findings of this study are openly available in https://github.com/mda-primetr/Spencer_et_al_2021. The R package (*asmbPLS*) implementing proposed method is available at https://github.com/RunzhiZ/asmbPLS.

## Consent for publication

Not applicable.

## Ethics approval and consent to participate

Not applicable.

## Supplementary Information

**Additional file 1**: The file contains the results for asmbPLS with different pre-defined quantile combinations and also the supplementary figures mentioned in the main text.

**Figure S1**. Prediction results for low dimension setting with lognormal distributed survival time and ***A*** = 0.5. **(A) *cr*** = 0.1; **(B) *cr*** = 0.3; **(C) *cr*** = 0.5; **(D) *cr*** = 0.7.

**Figure S2**. Prediction results for low dimension setting with Weibull distributed survival time and ***A*** = 2. **(A) *cr*** = 0.1; **(B) *cr*** = 0.3; **(C) *cr*** = 0.5; **(D) *cr*** = 0.7.

**Figure S3**. Prediction results for low dimension setting with Weibull distributed survival time and ***A*** = 0.5. **(A) *cr*** = 0.1; **(B) *cr*** = 0.3; **(C) *cr*** = 0.5; **(D) *cr*** = 0.7.

**Figure S4**. Sensitivity and specificity of the feature selection for low dimension setting with Weibull distributed survival time. **(A)** Sensitivity for microbiome block; **(B)** Sensitivity for metabolome block; **(C)** Specificity for microbiome block; **(D)** Specificity for metabolome block.

**Figure S5**. Prediction results for asmbPLS with different quantile combinations in low dimension setting with lognormal distributed survival time and ***A*** = 2. **(A) *cr*** = 0.1; **(B) *cr*** = 0.3; **(C) *cr*** = 0.5; **(D) *cr*** = 0.7.

**Figure S6**. Prediction results for asmbPLS with different quantile combinations in mixed dimension setting with lognormal distributed survival time and ***A*** = 2. **(A) *cr*** = 0.1; **(B) *cr*** = 0.3; **(C) *cr*** = 0.5; **(D) *cr*** = 0.7.

**Figure S7**. Prediction results for asmbPLS with different quantile combinations in high dimension setting with lognormal distributed survival time and ***A*** = 2. **(A) *cr*** = 0.1; **(B) *cr*** = 0.3; **(C) *cr*** = 0.5; **(D) *cr*** = 0.7.

**Figure S8**. Sensitivity and specificity of the feature selection for low dimension setting with lognormal distributed survival time for asmbPLS with different quantile combinations. **(A)** Sensitivity for microbiome block; **(B)** Sensitivity for metabolome block; **(C)** Specificity for microbiome block; **(D)** Specificity for metabolome block.

**Figure S9**. Sensitivity and specificity of the feature selection for mixed dimension setting with lognormal distributed survival time for asmbPLS with different quantile combinations. **(A)** Sensitivity for microbiome block; **(B)** Sensitivity for metabolome block; **(C)** Specificity for microbiome block; **(D)** Specificity for metabolome block.

**Figure S10**. Sensitivity and specificity of the feature selection for high dimension setting with lognormal distributed survival time for asmbPLS with different quantile combinations. **(A)** Sensitivity for microbiome block; **(B)** Sensitivity for metabolome block; **(C)** Specificity for microbiome block; **(D)** Specificity for metabolome block.

**Table S1**. Additional asmbPLS with different quantile combinations in different dimension settings.

