## Supplemental materials including the additional data analysis, supplemental tables, and supplemental figures. for "asmbPLS: Adaptive Sparse Multi-block Partial Least Square for Survival Prediction using Multi-Omics Data"

asmbPLS with different pre-defined quantile combinations

asmbPLS with the lowest quantile = 0.5 and asmbPLS with the lowest quantile = 0.9 were tested here. The details of the additional asmbPLS conducted are listed in **Table S1**. For simplicity, they were conducted in scenarios with lognormal distributed survival time only.

For asmbPLS with different quantile combinations (**Figure S5 - S7**), we found that the performances of different asmbPLS are similar except for the $\beta$ setting (4). In $\beta$ setting (4), since all the features have equal contributions to the outcome, the model with more features selected should have better performance, which heavily depends on the lowest quantile set in the quantile combinations. Therefore, asmbPLS with the lowest quantile = 0.5 shows the lowest MSE. For $\beta$ setting (1)(2)(3)(5)(6), since only a small proportion of the features are relevant, the performance of asmbPLS is affected more by the high quantile set in the quantile combinations. Therefore, asmbPLS with different quantile combinations show similar performance here due to the similar high quantile setting in the quantile combinations.

In addition, as we expected, among asmbPLS with different quantile combinations (**Figure S8 - S10**), asmbPLS with higher quantile combinations tend to have lower sensitivity and higher specificity regardless of the scenarios.

**Supplementary Figures**

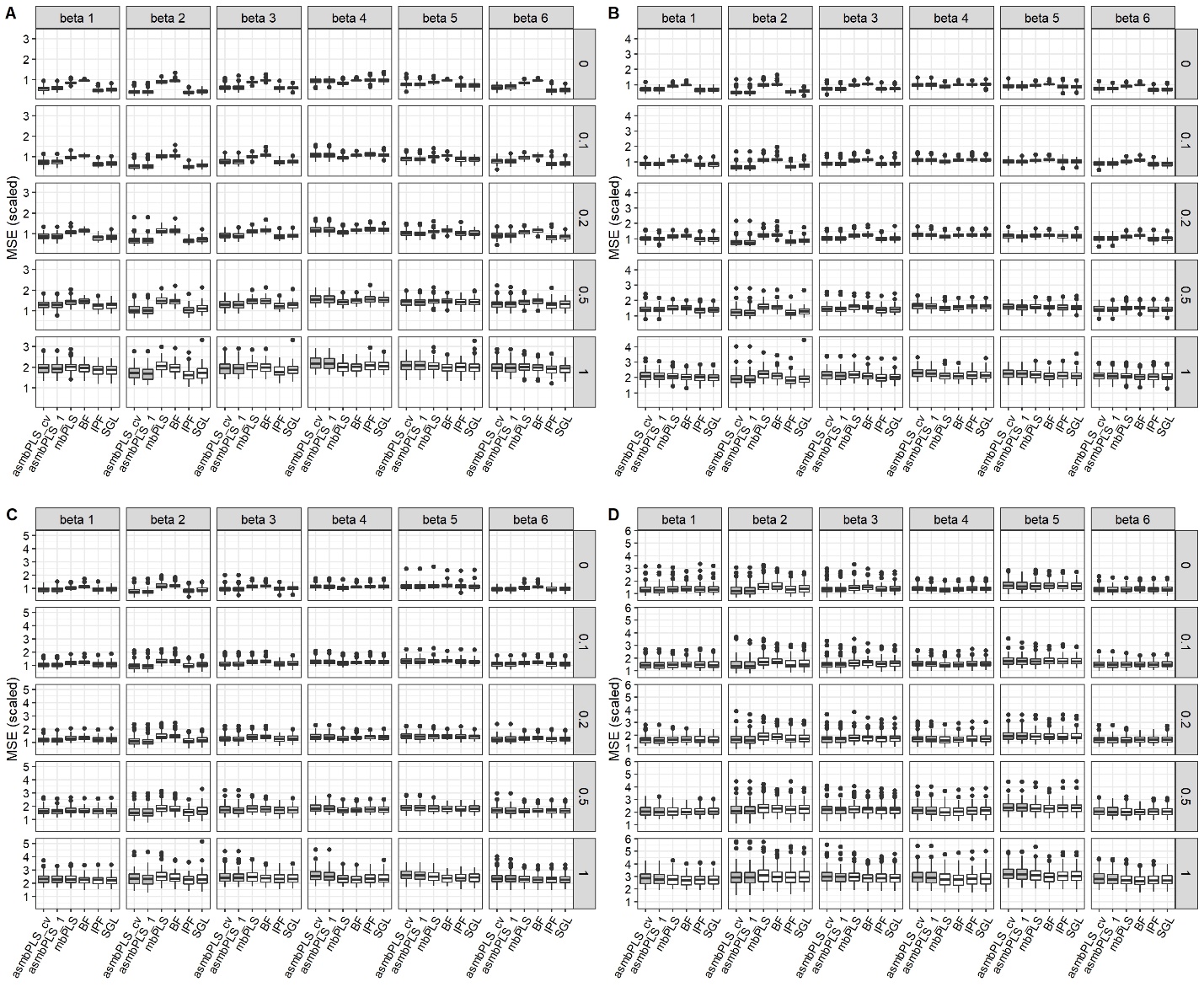

**Figure S1**. Prediction results for low dimension setting with lognormal distributed survival time and $A$ = 0.5. **(A)** $cr$ = 0.1; **(B)** $cr$ = 0.3; **(C)** $cr$ = 0.5; **(D)** $cr$ = 0.7.

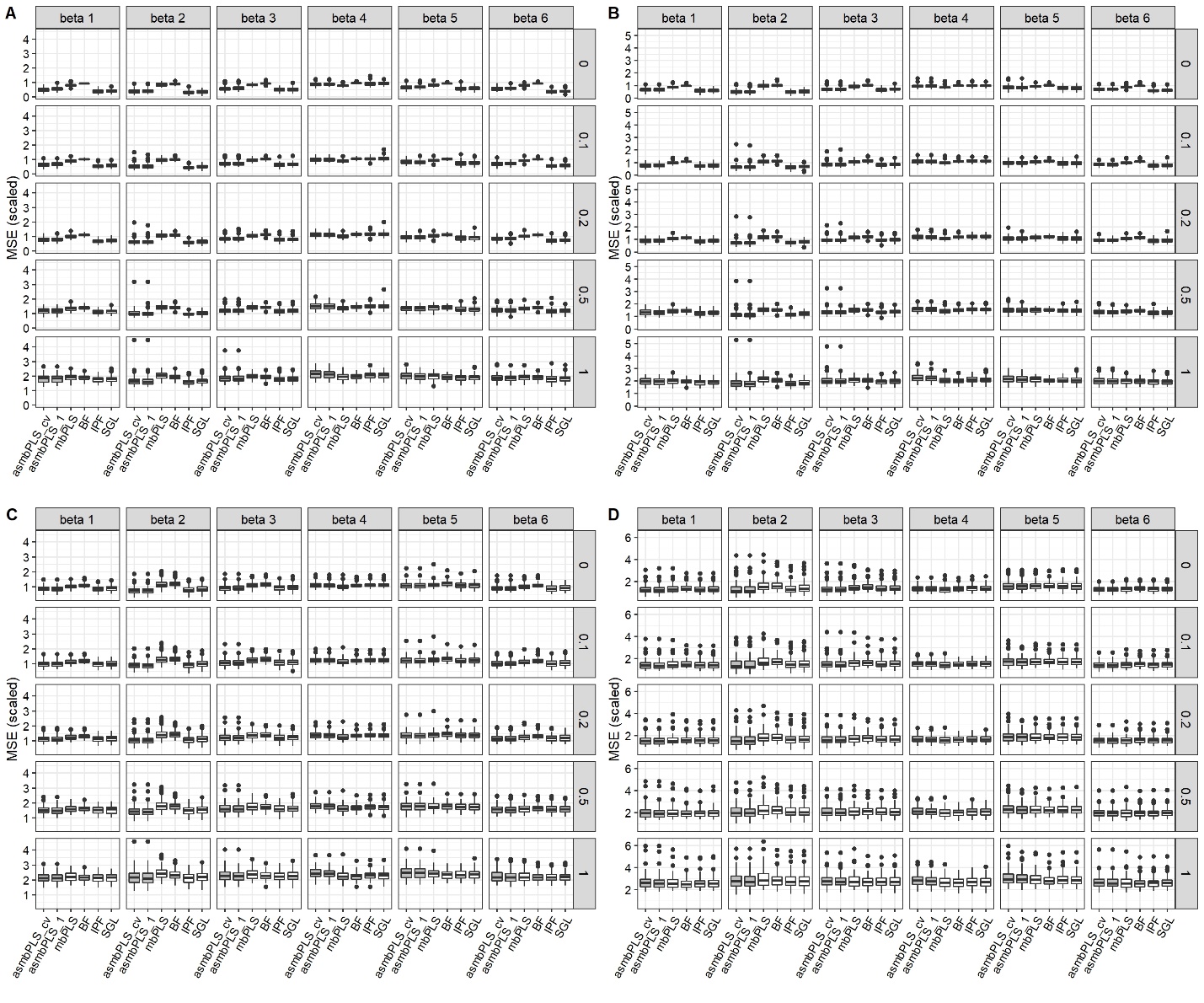

**Figure S2**. Prediction results for low dimension setting with Weibull distributed survival time and $A$ = 2. **(A)** $cr$ = 0.1; **(B)** $cr$ = 0.3; **(C)** $cr$ = 0.5; **(D)** $cr$ = 0.7.

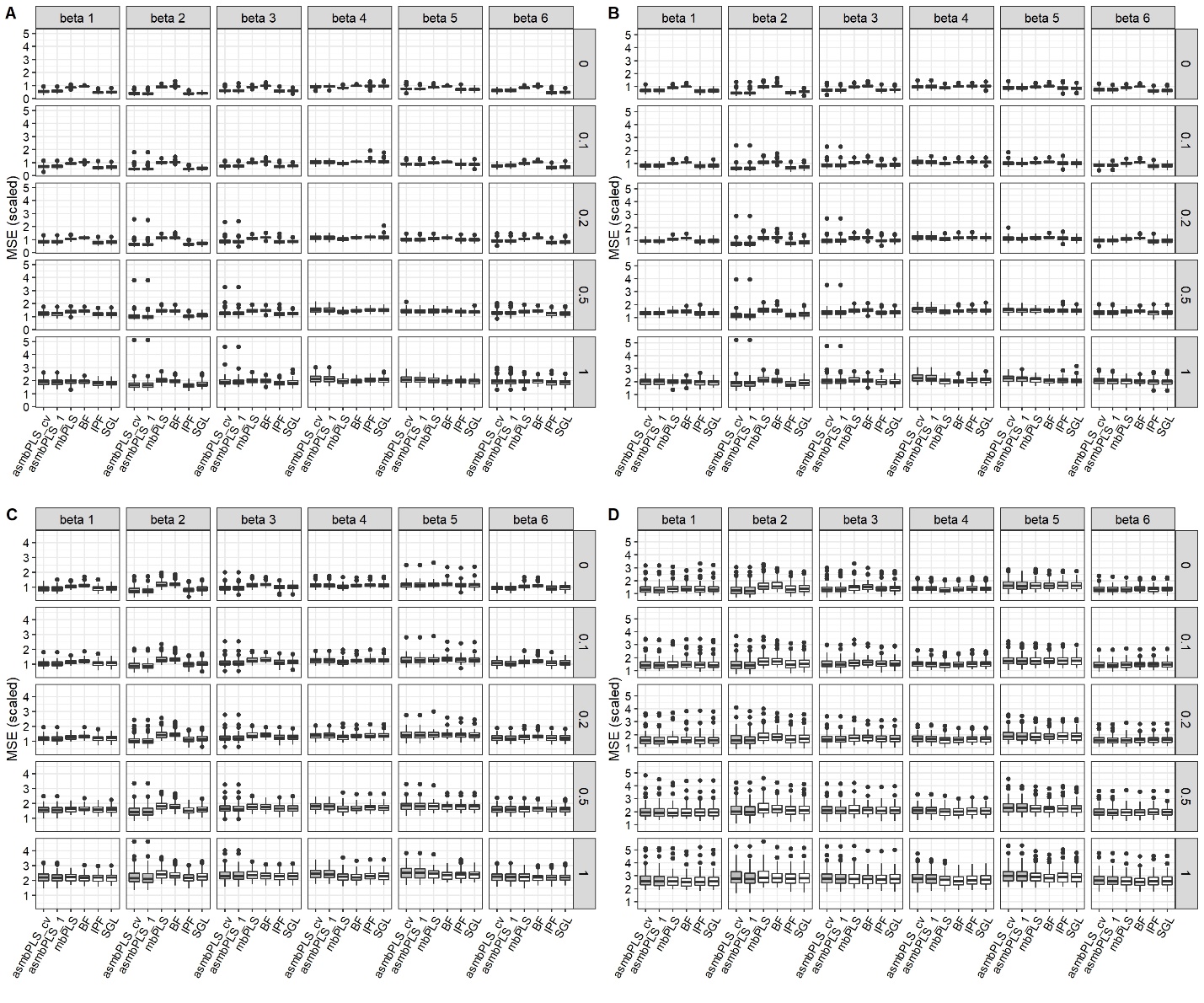

**Figure S3**. Prediction results for low dimension setting with Weibull distributed survival time and $A$ = 0.5. **(A)** $cr$ = 0.1; **(B)** $cr$ = 0.3; **(C)** $cr$ = 0.5; **(D)** $cr$ = 0.7.

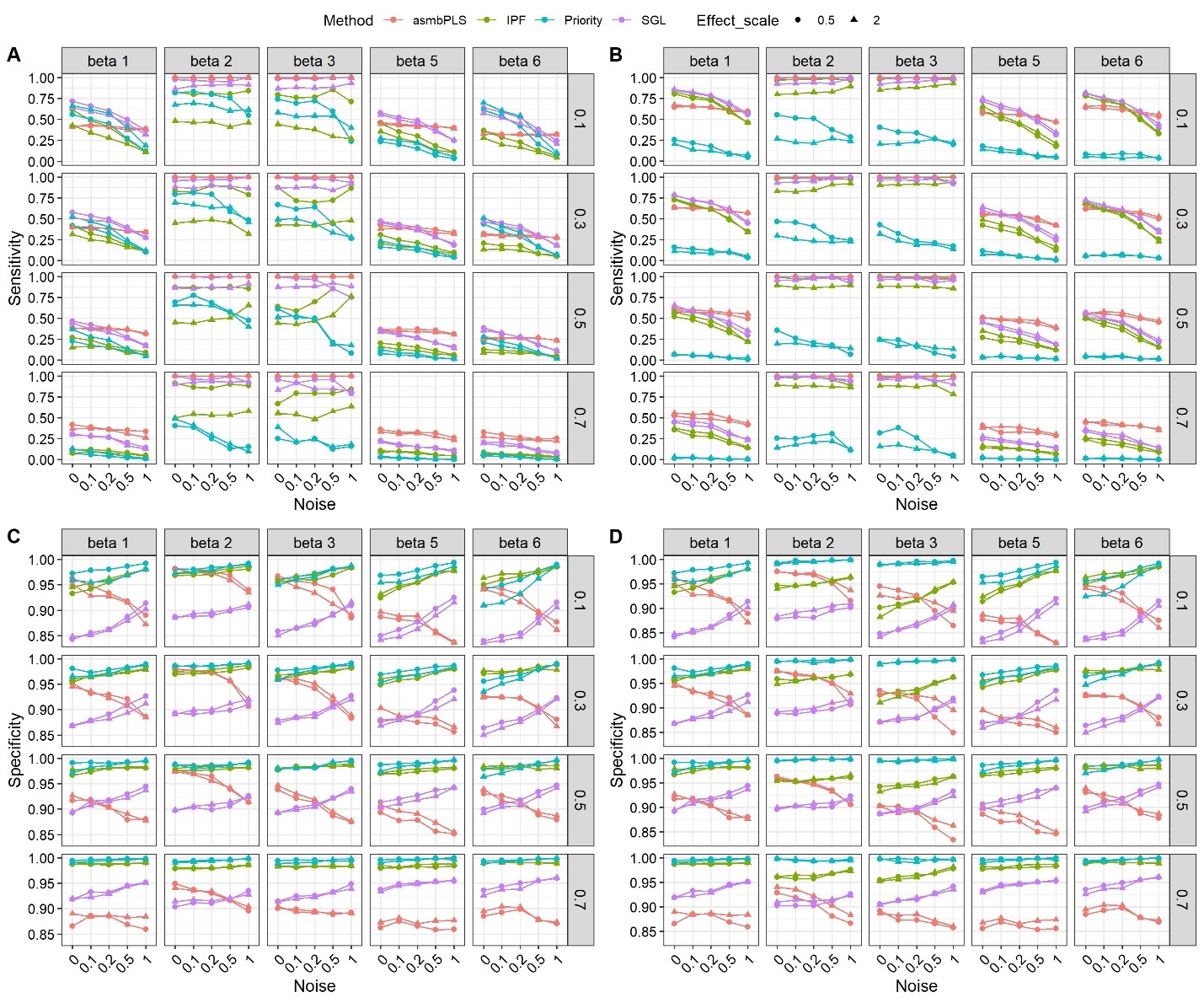

**Figure S4.** Sensitivity and specificity of the feature selection for low dimension setting with Weibull distributed survival time. **(A)** Sensitivity for microbiome block; **(B)** Sensitivity for metabolome block; **(C)** Specificity for microbiome block; **(D)** Specificity for metabolome block.

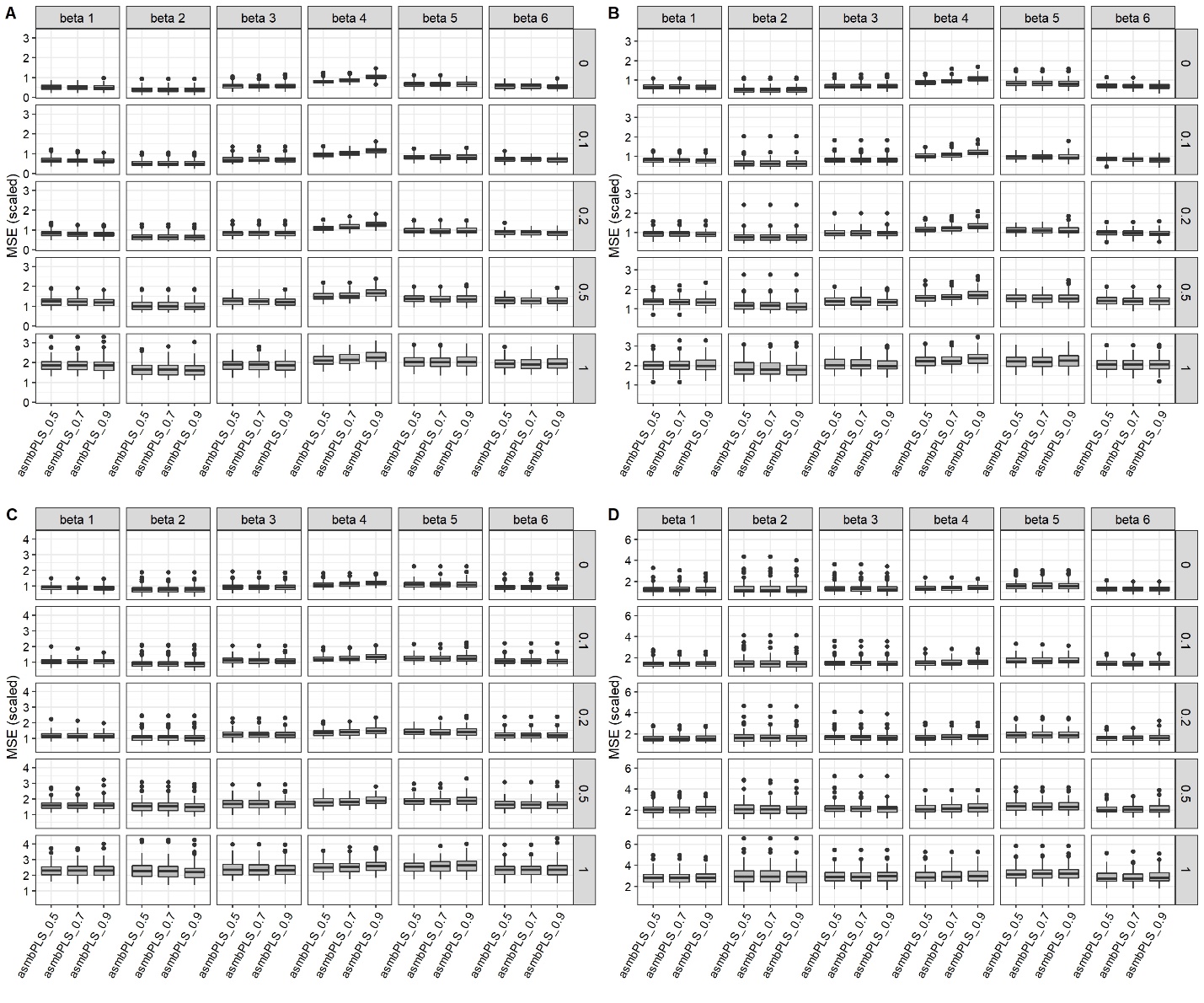

**Figure S5**. Prediction results for asmbPLS with different quantile combinations in low dimension setting with lognormal distributed survival time and $A$ = 2. **(A)** $cr$ = 0.1; **(B)** $cr$ = 0.3; **(C)** $cr$ = 0.5; **(D)** $cr$ = 0.7.

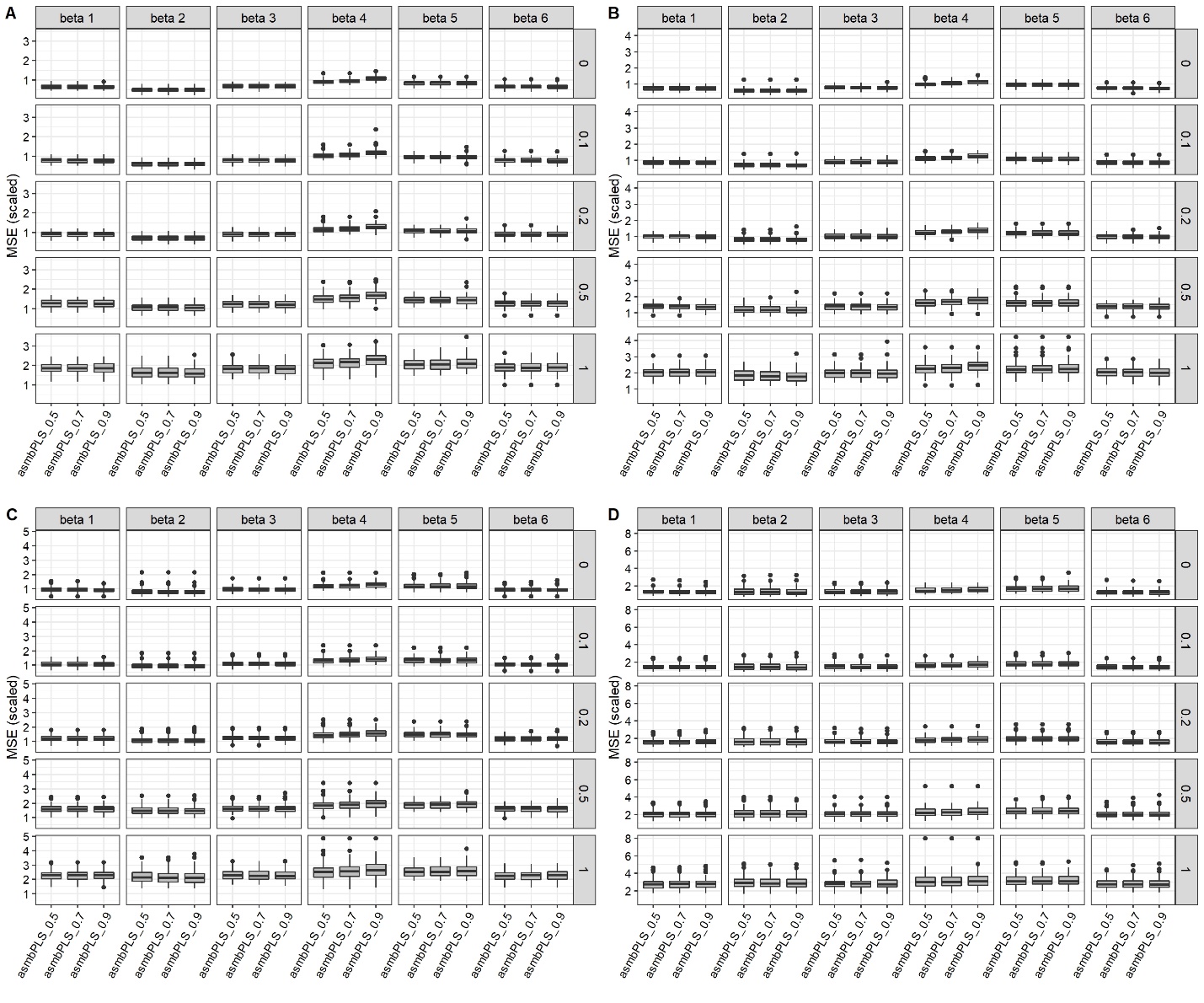

**Figure S6**. Prediction results for asmbPLS with different quantile combinations in mixed dimension setting with lognormal distributed survival time and $A$ = 2. **(A)** $cr$ = 0.1; **(B)** $cr$ = 0.3; **(C)** $cr$ = 0.5; **(D)** $cr$ = 0.7.

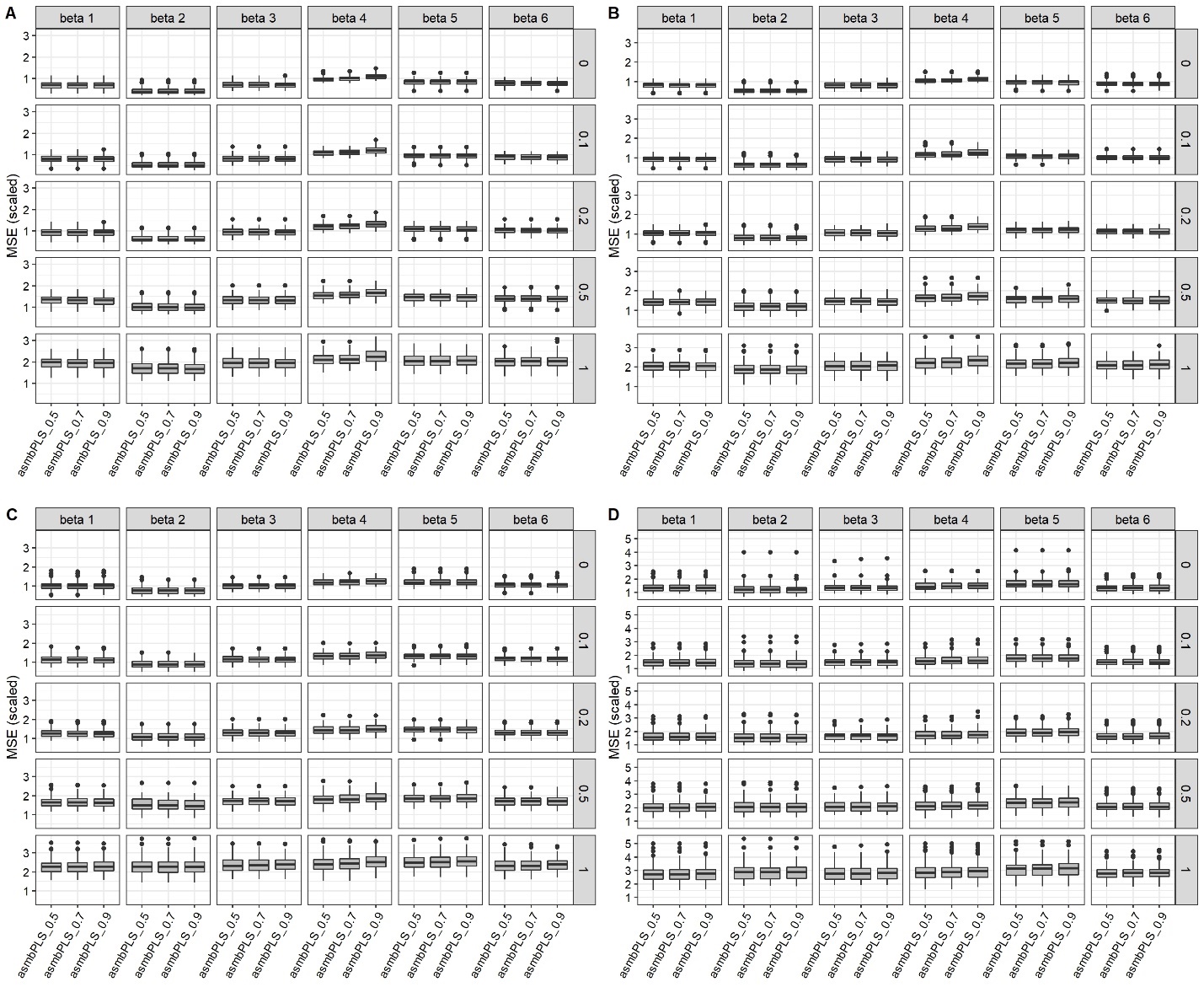

**Figure S7**. Prediction results for asmbPLS with different quantile combinations in high dimension setting with lognormal distributed survival time and $A$ = 2. **(A)** $cr$ = 0.1; **(B)** $cr$ = 0.3; **(C)** $cr$ = 0.5; **(D)** $cr$ = 0.7.

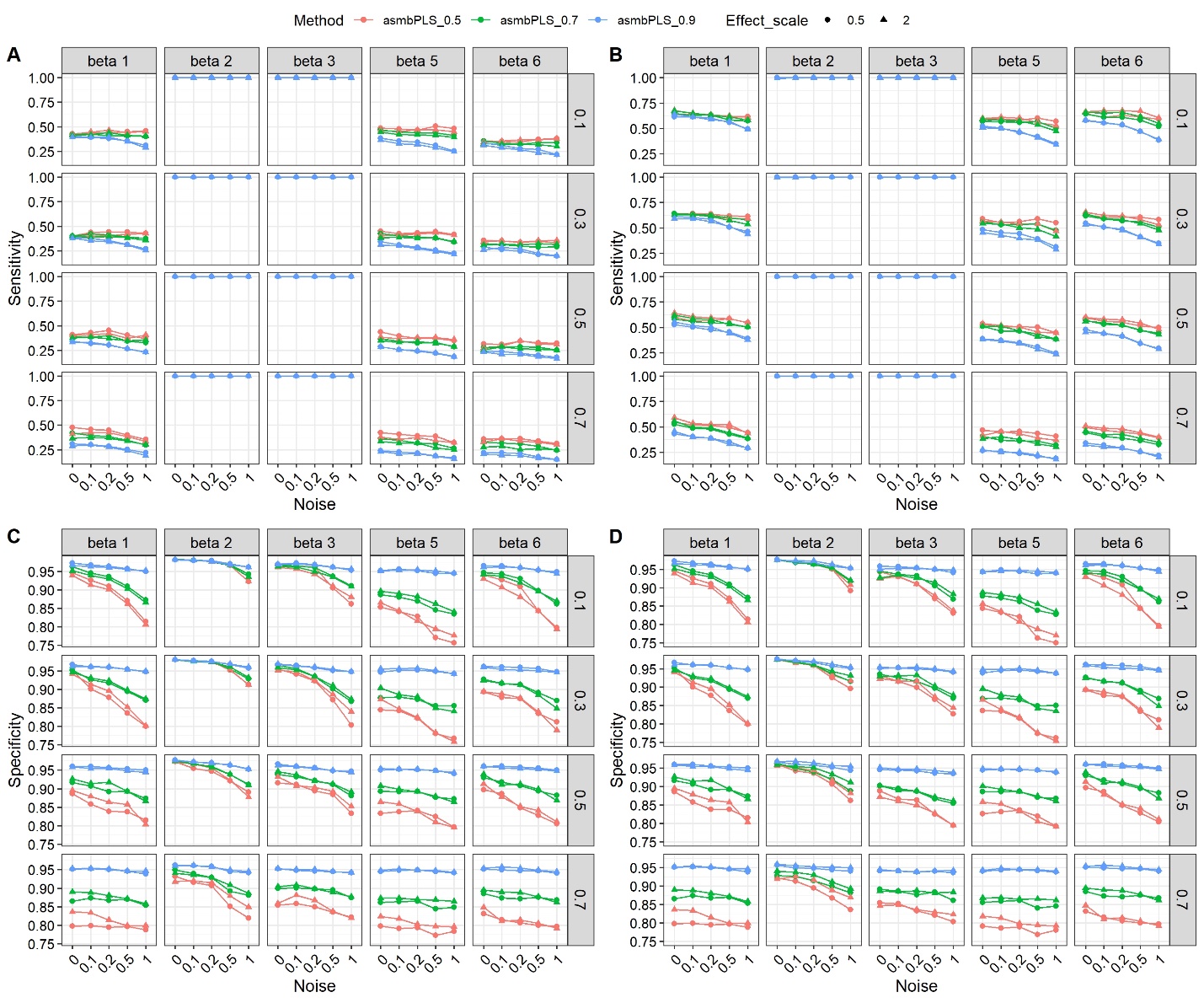

**Figure S8.** Sensitivity and specificity of the feature selection for low dimension setting with lognormal distributed survival time for asmbPLS with different quantile combinations. **(A)** Sensitivity for microbiome block; **(B)** Sensitivity for metabolome block; **(C)** Specificity for microbiome block; **(D)** Specificity for metabolome block.

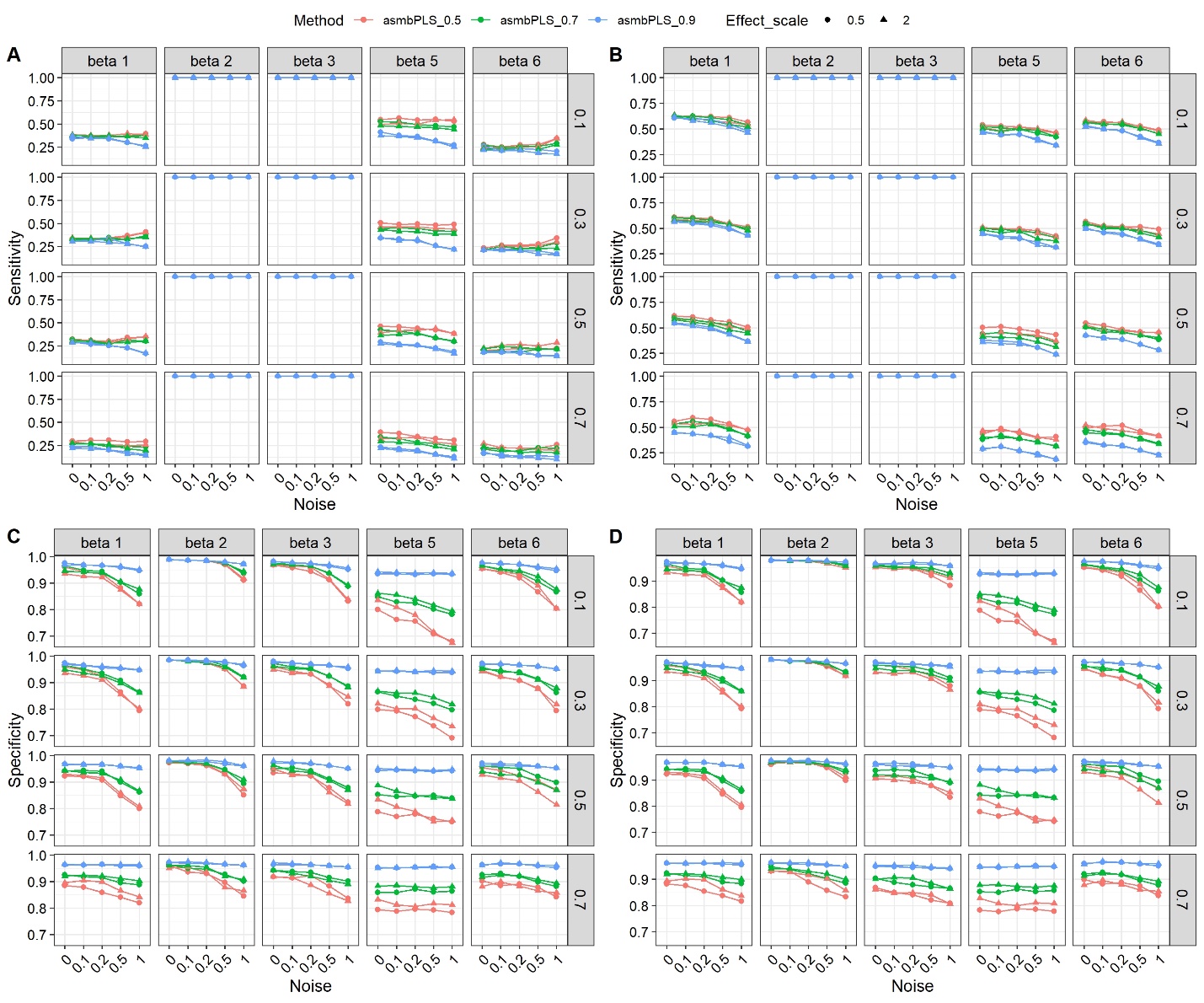

**Figure S9.** Sensitivity and specificity of the feature selection for mixed dimension setting with lognormal distributed survival time for asmbPLS with different quantile combinations. **(A)** Sensitivity for microbiome block; **(B)** Sensitivity for metabolome block; **(C)** Specificity for microbiome block; **(D)** Specificity for metabolome block.

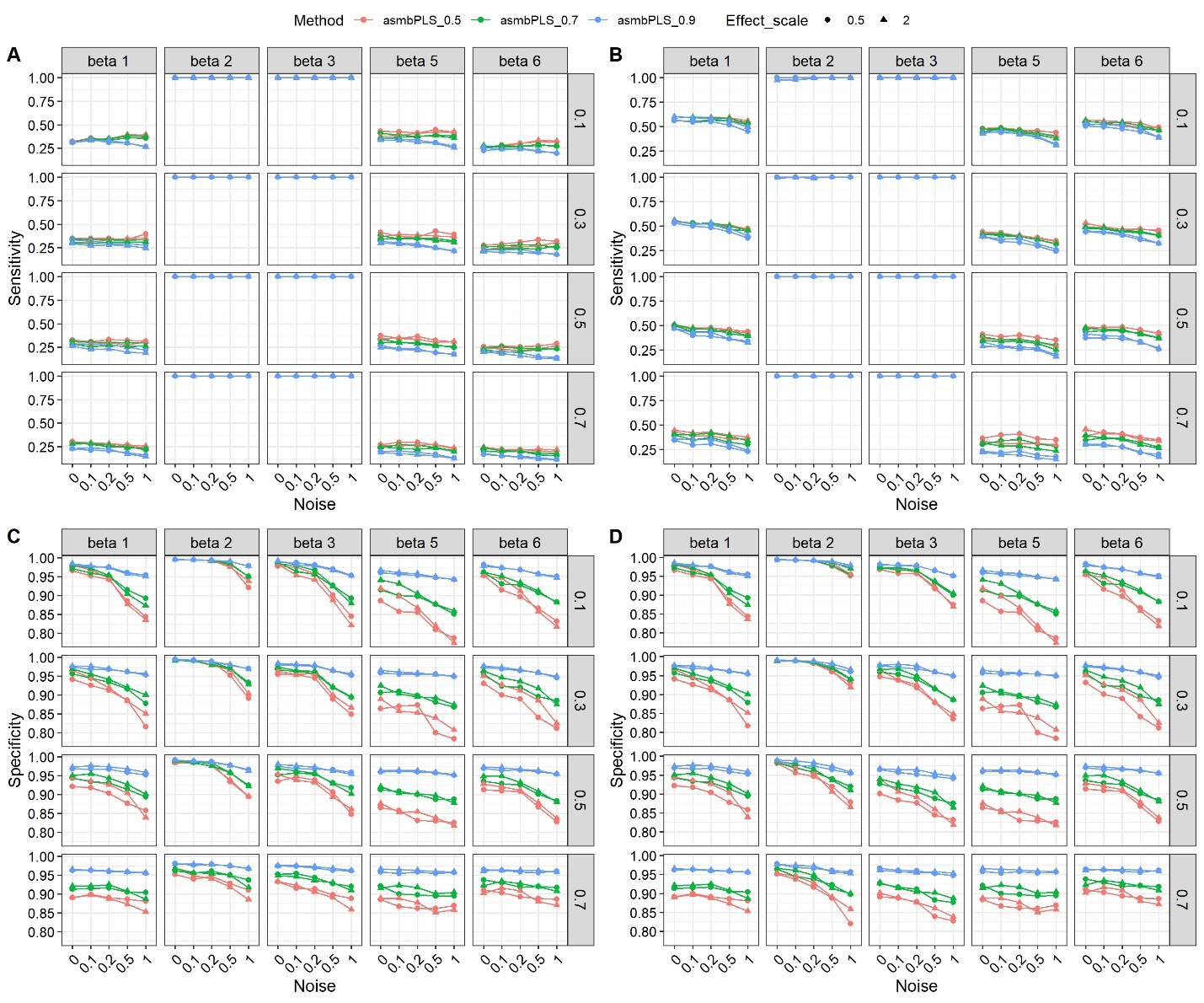

**Figure S10.** Sensitivity and specificity of the feature selection for high dimension setting with lognormal distributed survival time for asmbPLS with different quantile combinations. **(A)** Sensitivity for microbiome block; **(B)** Sensitivity for metabolome block; **(C)** Specificity for microbiome block; **(D)** Specificity for metabolome block.

**Supplementary Tables**

**Table S1**. Additional asmbPLS with different quantile combinations in different dimension settings.

| Dimension | $\mathrm{quantil}e_{block_{1}}$ | $\mathrm{quantil}e_{block_{2}}$ |
| --- | --- | --- |
| Low | {0.5, 0.6, 0.7, 0.8, 0.9, 0.95, 0.975} | {0.5, 0.6, 0.7, 0.8, 0.9, 0.95, 0.975} |
|  | {0.9, 0.925, 0.95, 0.975} | {0.9, 0.925, 0.95, 0.975} |
| Mixed | {0.5, 0.6, 0.7, 0.8, 0.9, 0.95, 0.975, 0.99, 0.995} | {0.5, 0.6, 0.7, 0.8, 0.9, 0.95, 0.975} |
|  | {0.9, 0.925, 0.95, 0.975, 0.99, 0.995} | {0.9, 0.925, 0.95, 0.975} |
| High | {0.5, 0.6, 0.7, 0.8, 0.9, 0.95, 0.975, 0.99, 0.995} | {0.5, 0.6, 0.7, 0.8, 0.9, 0.95, 0.975, 0.99, 0.995} |
|  | {0.9, 0.925, 0.95, 0.975, 0.99, 0.995} | {0.9, 0.925, 0.95, 0.975, 0.99, 0.995} |
